# DNA damage drives antigen diversification through mosaic Variant Surface Glycoprotein (VSG) formation in *Trypanosoma brucei*

**DOI:** 10.1101/2024.03.22.582209

**Authors:** Jaclyn E. Smith, Kevin J. Wang, Erin M. Kennedy, Jill M.C. Hakim, Jaime So, Alexander K. Beaver, Aishwarya Magesh, Shane D. Gilligan-Steinberg, Jessica Zheng, Bailin Zhang, Dharani Narayan Moorthy, Elgin Henry Akin, Lusajo Mwakibete, Monica R. Mugnier

**Author notes:** Correspondence (M.R.M.).

## Abstract

Antigenic variation, using large genomic repertoires of antigen-encoding genes, allows pathogens to evade host antibody. Many pathogens, including the African trypanosome *Trypanosoma brucei,* extend their antigenic repertoire through genomic diversification. While evidence suggests that *T. brucei* depends on the generation of new variant surface glycoprotein (VSG) genes to maintain a chronic infection, a lack of experimentally tractable tools for studying this process has obscured its underlying mechanisms. Here, we present a highly sensitive targeted sequencing approach for measuring VSG diversification. Using this method, we demonstrate that a Cas9-induced DNA double-strand break within the VSG coding sequence can induce VSG recombination with patterns identical to those observed during infection. These newly generated VSGs are antigenically distinct from parental clones and thus capable of facilitating immune evasion. Together, these results provide insight into the mechanisms of VSG diversification and an experimental framework for studying the evolution of antigen repertoires in pathogenic microbes.

## Introduction

Pathogen survival in a host depends upon effective and continuous immune evasion. Several bacteria and eukaryotic pathogens have adopted the strategy of antigenic variation to evade host immunity, a process in which they continuously alter antigenic surface proteins to escape the host’s adaptive immune response. The African trypanosome *Trypanosoma brucei*, a unicellular eukaryotic parasite and causative agent of human and animal African trypanosomiasis, uses an especially sophisticated system of antigenic variation. The parasite, which remains extracellular throughout infection and thus faces a perpetual onslaught of host antibody, periodically “switches” expression of a surface coat consisting of 10^7^ copies of a single, immunogenic protein known as the Variant Surface Glycoprotein (VSG). This process allows parasites to escape host antibody and maintain a chronic infection.

Although the *T. brucei* VSG repertoire contains thousands of VSGs, this repertoire is probably too small to maintain a chronic infection through VSG switching alone. During an infection, each *T. brucei* parasite expresses a single VSG at a time from one of ∼15 telomeric Bloodstream Expression Sites (the “active” BES)^1^. The remaining VSG-encoding genes are stored in other expression sites, subtelomeric arrays, and minichromosomes, all of which remain transcriptionally silenced^2^. The parasite switches its VSG either by transcriptional activation of a silent BES (*in situ* switching) or through a gene conversion event in which a new VSG is copied into the active expression site. While gene conversion-based switching allows for the activation of VSGs outside of a BES, analysis of the *T. brucei* genome has shown that only ∼20% of the VSGs in the parasite genome are full-length genes encoding a functional VSG protein. The remaining ∼80% of VSGs in the parasite genome consist of pseudogenes or gene fragments^2^ and cannot immediately be used for immune evasion through *in situ* or gene conversion switching. Moreover, the number of VSGs expressed in a population of parasites at a single time during experimental infection sometimes exceeds the total number of intact VSGs in the parasite genome^3,4^, further indicating that the repertoire of intact VSGs is insufficient to achieve the antigenic diversity required to maintain a chronic infection.

Evidence suggests that *T. brucei* deals with this shortage of antigens through diversification of the VSG repertoire. Many studies of experimental infections in mice have shown that novel VSGs, generated during infection, predominate at later stages of infection^3,5,6^, while analysis of parasites from natural human infections revealed expressed VSGs that were nearly completely absent from the genomes of contemporary field isolates^7^. These observations suggest that the generation of new VSGs plays a critical role in sustaining *T. brucei* antigenic variation.

There are two mechanisms thought to be responsible for extending the VSG repertoire: mosaic formation and *de novo* point mutation. Mosaic VSGs form when two or more VSG genes combine through segmental gene conversion to form a novel VSG. This mechanism allows parasites to access pseudogenes and VSG fragments within the repertoire. While mosaic VSGs have been observed in the literature, they are often found under extreme experimental conditions^8–12^ or late during infection^3,5,13,14^ making it difficult to discern how exactly they arose. VSGs also appear to occasionally acquire *de novo* point mutations, though these can be difficult to distinguish from a small gene conversion event using a similar donor VSG. The origins of these mutations also remain mysterious, though newly acquired mutations seem rare^9,15–17^. Ultimately, *de novo* mutation of VSGs would allow parasites to generate new VSG sequences regardless of the contents of their repertoire, further amplifying diversity.

Despite its clear importance, the mechanisms driving VSG diversification, whether by mutation or recombination, remain poorly understood. It is plausible that DNA damage and repair may play a role in either mechanism, as expressed VSG genes sit between two highly repetitive and damage-prone stretches of DNA, the conserved 70bp repeat and the telomere^2,18^. DNA breaks are frequently observed within the 70bp repeats at the active expression site^19,20^ and near telomeres within silent BESs^20^. Experimental evidence implicates DNA damage in VSG switching more generally, as a DNA double-strand break induced upstream of the VSG induces a switch^19,20^, but it is unknown whether DNA damage could also generate new VSGs.

There is a dearth of experimental evidence for the mechanisms driving VSG diversification because there has been no controlled and reproducible *in vitro* system for studying the process. Instead, researchers have had to rely on observation alone, characterizing the sequences of expressed VSGs that were isolated by chance. This approach has been crucial for generating hypotheses but is insufficient for deciphering the mechanisms driving the process. Here, we present a comprehensive toolkit for the controlled and reproducible study of the diversification of individual VSGs. Using a highly sensitive barcode-based targeted RNA-seq approach, we show that DNA double-strand breaks can trigger the formation of mosaic VSGs that are identical to those observed *in vivo* during infection. Homology appears to drive donor VSG selection, with microhomology between the parent and donor VSG flanking the site of recombination in almost every event. We demonstrate that homologous donor VSGs can be utilized as a template for DNA repair regardless of their genomic location. Finally, we observe that break-induced diversification is most efficient in the portion of the VSG gene encoding the exposed top lobe of the VSG N-terminal domain, allowing very small changes in the VSG protein to facilitate large changes in antibody binding. We suggest that this could represent a potential hypervariable region within the VSG, facilitating diversification in the region of the VSG protein where host antibody is most likely to bind.

## Results

### DNA double-strand breaks trigger the formation of mosaic VSGs

A number of studies have suggested that VSG diversification occurs, at least some of the time, within the active expression site^3,16^, with the sequence of the actively expressed VSG being altered through recombination and/or mutation. For this reason, we focused our analyses on diversification of the actively expressed VSG. Because a double-strand DNA break upstream of the VSG within the active BES is known to induce switching^19,20^, we hypothesized that a break within the VSG coding sequence might result in mosaic VSG formation. To investigate this, we engineered tetracycline-inducible Cas9 expressing EATRO1125 parasites, which express the VSG AnTat1.1, and induced breaks across the AnTat1.1 coding sequence using a set of guide RNAs.

To evaluate potential recombination outcomes, we developed VSG Anchored Multiplex PCR Sequencing (VSG-AMP-seq), a technique that overcomes previous obstacles to studying VSG diversification by providing both high-throughput and highly accurate sequences (1A). Our approach uses long unique molecular indexes (UMIs) to generate high-confidence consensus sequences^21^. Consolidating reads into consensus sequences allows errors like PCR chimeras, which occur during later cycles of PCR and therefore represent a minority of sequences in a consensus group^22^, to be eliminated while true events are retained. Libraries are prepared by fragmenting VSG-specific cDNA. Fragments are then end-repaired, A-tailed and ligated to universal adapters containing a 25bp UMI (1A). A VSG target of interest is selected, and a series of staggered primers are designed to cover the length of the target VSG’s coding sequence (1B). By pairing target-specific primers with a universal reverse adapter primer, target VSG fragments are amplified within a sample regardless of their identity. Mosaic reads are defined as those for whom a portion of the read matches the target and the remainder matches another VSG (the “donor VSG”) within the VSG repertoire (VSGnome) of the strain being studied^2^. Due to its selective, target-specific amplification, this method can sensitively detect thousands of rare diversification events, even from mixed samples where diversified VSGs are a minority of the population. To validate this approach, we mixed together at a known ratio two parasite clones expressing two distinct VSGs that shared sufficient sequence similarity to make them prone to the generation of PCR chimeras. We then performed VSG-AMP-seq on RNA extracted from this parasite mixture. This analysis revealed very few erroneous recombination events, even without UMI consolidation, demonstrating that VSG-AMP-seq is highly accurate and detects true mosaic recombination events (Supplemental Figure 1).

**Figure 1.**
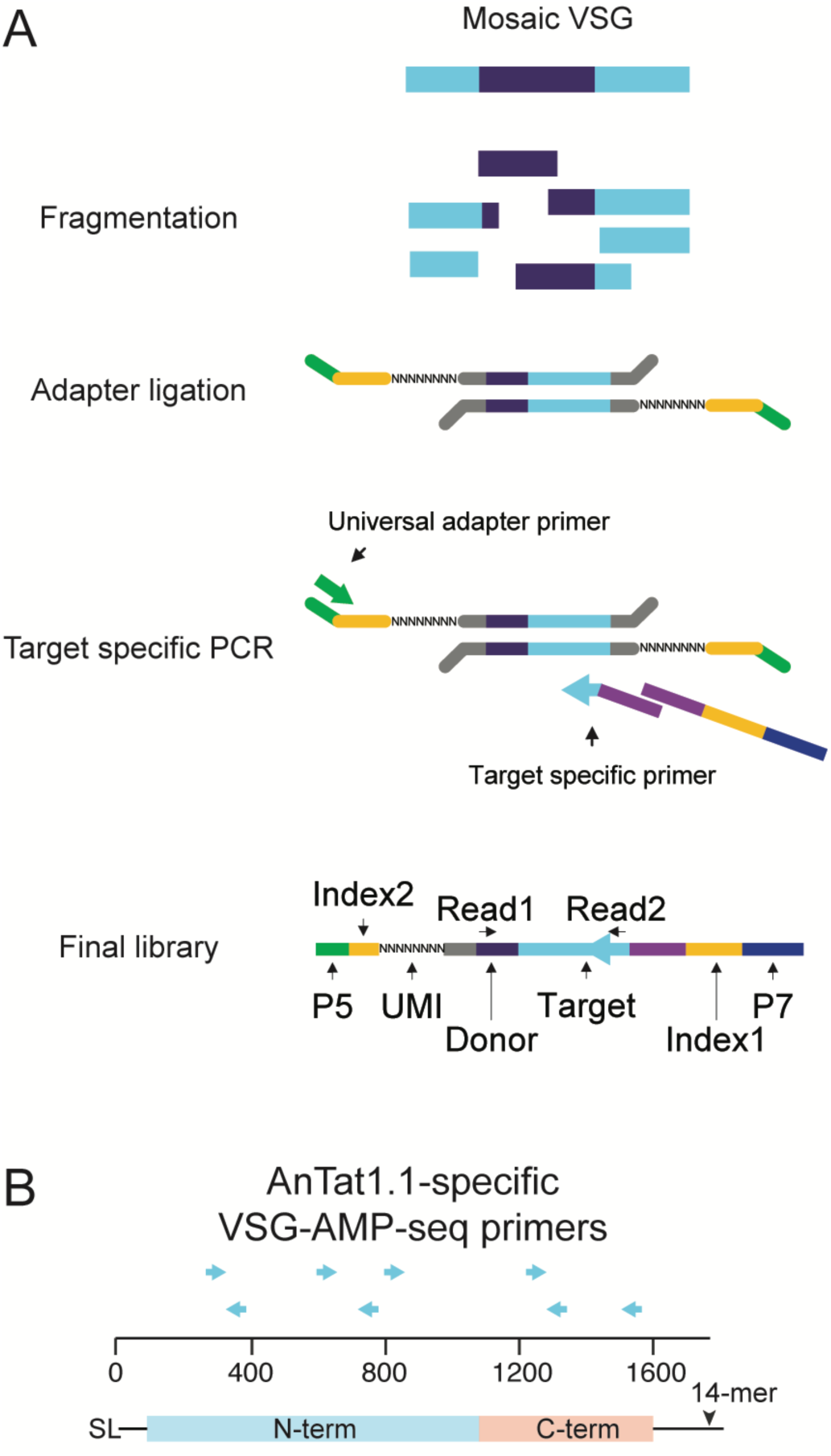
VSG-AMP-seq protocol. A) Schematic of the library prep for VSG-AMP-seq. Figure adapted from ^65^. B) Locations of target specific primers for VSG AnTat1.1. SL = 5′ splice leader sequence, 14-mer = 3′ sequence conserved in all VSG transcripts

To induce breaks across AnTat1.1, we induced Cas9 expression for 24 hours and then transfected in DNA amplicons containing a T7 promoter and a guide RNA targeting various regions throughout the AnTat1.1 coding sequence to induce breaks in the VSG. Parasites were collected two days after transfection of the guide, and mosaic derivatives of AnTat1.1 were analyzed by VSG-AMP-seq (2A; Supplementary Figures 2A, 2B & 3C). We detected thousands of recombination events from two independently generated Cas9 clones (C1 = 5956, C2 = 4488).

**Figure 2.**
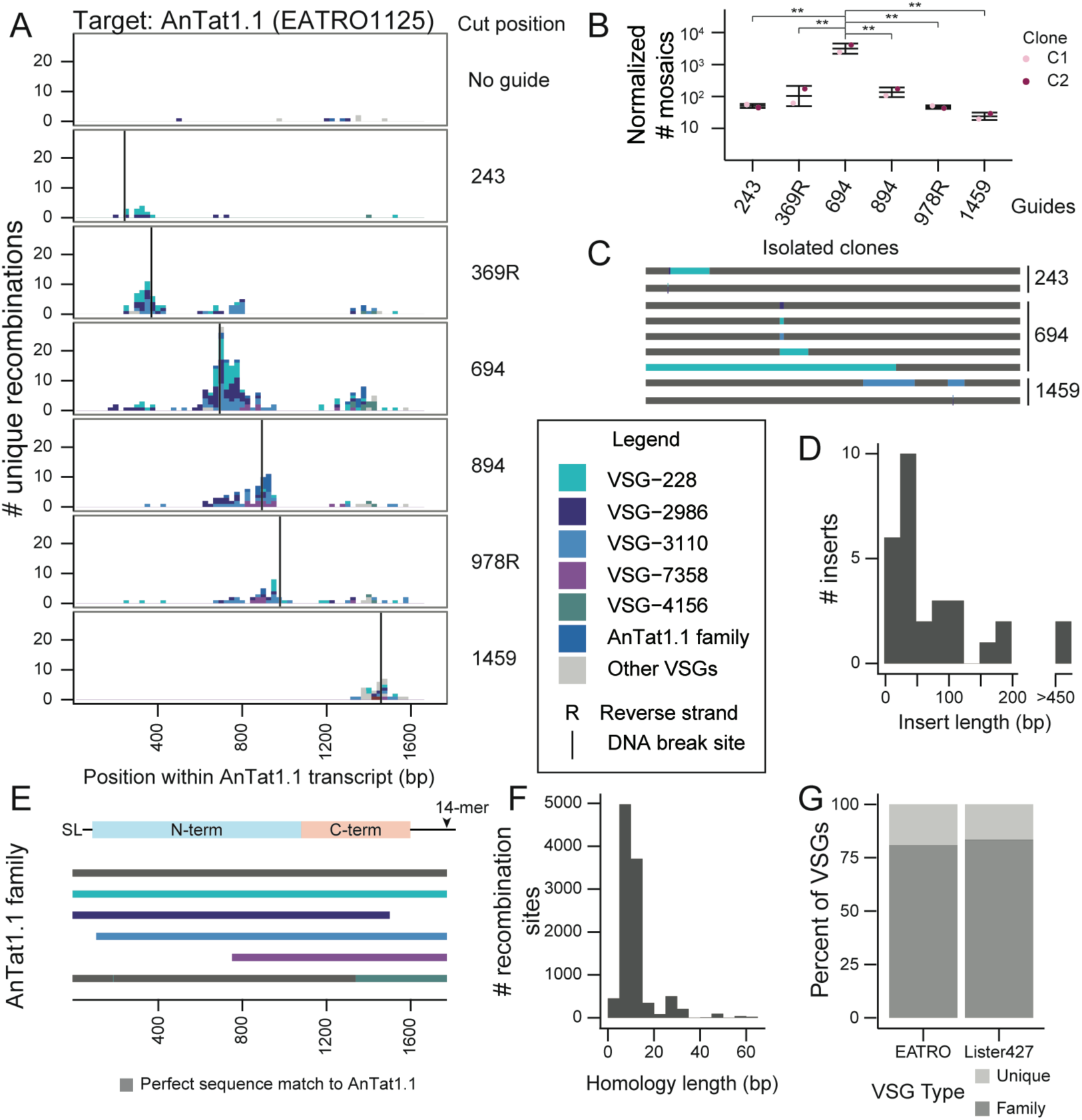
DNA double-strand breaks trigger mosaic VSG formation when homology is available. A) A histogram of unique recombination events identified along the AnTat1.1 transcript. The Cas9 DNA break site is indicated by a vertical line. Cut positions, relative to the 5’ end of the VSG transcript are: 243, 369, 694, 894, 978, and 1459. Sections of the histogram are colored to indicate donor VSG identified. R indicates a guide that binds to the reverse strand. The midpoint of the perfect homology between AnTat1.1 and the donor VSG at the recombination site is plotted. If a mosaic sequence matched >1 potential donor VSG, the average recombination position was plotted. B) Quantification of mosaic recombination events induced by DNA breaks. The number of recombination events detected within 250bp up or downstream of the cut site was normalized to the number of total unanchored reads aligning within that region compared to the unanchored read count from the region with the smallest coverage to control for sequencing depth. (n=2, two independent clones) Statistical significance was determined with a one-way ANOVA with post-hoc Tukey HSD (** p <0.01) Mean ± s.d. C) Schematics of mosaic VSGs from parasite clones isolated after DNA breaks within AnTat1.1. Representative sequences shown. D) A histogram of donor VSG insertion lengths identified in all isolated mosaic-expressing clones. The insert length only includes newly inserted sequence and does not include recombination sites. E) A schematic of the AnTat1.1 family aligned to the AnTat1.1 transcript. Gray sequences are a perfect match to AnTat1.1. F) A histogram of the length of the shared identity between AnTat1.1 and the donor VSG at each recombination site. G) Quantification of types of VSGs within the VSGnomes from EATRO1125 and Lister427 parasites. The Lister427 VSGnome has 5 VSGs which are perfectly duplicated without any other family members. SL = 5′ splice leader sequence, 14-mer = 3′ sequence conserved in all VSG transcripts

Our analysis revealed diverse mosaic recombination events centered around each break site. Such events were virtually absent from the negative (no guide) control, suggesting this process does not occur at a high rate in the absence of a trigger or in the presence of Cas9 alone. Notably, as the DNA breaks progressed further away from the center of the AnTat1.1 coding sequence, the frequency of mosaic formation decreased dramatically (2B). This did not appear to be related to the guide sequences (Supplemental Figure 2C-G) or to guide cutting efficiency (Supplemental Figure 2H). To ensure that the observed mosaic events represented mosaic VSGs that were truly expressed by the parasite, and not technical artifacts or VSGs incapable of being stably expressed by *T. brucei*, we also obtained individual clones of parasites expressing AnTat1.1-derived mosaic VSGs. Using parasite lines that stably express a VSG-targeted guide RNA, we induced Cas9 expression for 7 days (2C; Supplemental Figure 2A). We isolated 29 mosaic-expressing parasite clones, 4 from guide 243, 22 from guide 694, and 3 from guide 1459, from a total of 5 parental cell lines. These parasites expressed VSGs containing recombination events identical to those observed using VSG-AMP-seq. These results suggest that DNA damage within the active VSG can trigger the formation of mosaic VSGs, with recombination events centered around the site of DNA damage.

### Homology drives mosaic VSG formation

Analysis of the mosaic recombination events detected by VSG-AMP-seq revealed an important role for sequence homology in the formation of mosaic VSGs. Within the isolated mosaic clones, we observed short insertions (< 200 bp, average = 49 bp, median = 23 bp) predominating among the events (2D). Although read lengths limited our ability to detect larger insertions with VSG-AMP-seq, approximately 55-60% of the mosaic recombination reads detected by VSG-AMP-seq contained the same short insertions (average C1 = 46 bp, average C2 = 45.8 bp) (Supplemental Figure 3C). Almost all (C1= 99.71%, C2 = 99.46%) of these recombination events occurred within a region of shared sequence between AnTat1.1 and each donor VSG, with an average length of ∼9 bps (average C1 = 9.13 bp, average C2 = 9.44 bp, median = 6 bp) (2F; Supplemental Figure 1A). Notably, only a small number of donor VSGs were used for the majority of recombination events. Upon closer inspection, these donors are members of a 6-VSG family that contains AnTat1.1 and represent the only sequences within the EATRO1125 VSGnome with significant homology to this VSG (2E). N-terminal recombination events appear restricted to just these family members (C1 = 100%, C2 = 99.87%). Together, these results suggest that VSG sequence homology influences the outcome of VSG recombination after a DNA break.

Not all VSGs share sequence homology with other members of the VSG repertoire, however. To determine the outcome following DNA damage of a VSG when there are no homologous donors available, we used the same Cas9 system in the commonly used Lister427 *T. brucei* line to cut the actively expressed VSG, VSG-2, which lacks homologous family members. After inserting stably expressed guides targeting VSG-2 into the genome, we induced Cas9 expression for 7 days. Parasite clones isolated after a break in VSG-2 no longer expressed VSG-2, and the VSGs expressed by these clones showed no evidence of mosaic recombination (Supplemental Figure 3B; cut position 707, n = 4; cut position 1082, n = 3). This further supports the hypothesis that sequence homology between the parent and donor VSG is required for mosaic VSG formation.

We sought to determine what proportion of the VSG repertoire contains VSGs that are members of VSG families and thus capable of diversifying through break-induced mosaic formation. We performed an all vs all BLAST to cluster VSGs based upon sequence similarity within both the EATRO1125 and Lister427 parasite strains and found that most (>75%) of the known VSG sequences are capable of diversifying through the mechanisms described here (2G). We supplemented this analysis with an orthogonal clustering method and estimated a similar proportion of diversifiable VSGs (Supplemental Figure 3A).

### Donor VSGs act as templates for break-induced VSG diversification

Although most AnTat1.1-derived mosaics were identical to the putative donor VSG, many of these insertions altered only a few bps in AnTat1.1. We thus reasoned that it was possible that these events were *de novo* mutations created during DNA repair that happened to match the putative genome-encoded donor VSG. To test this possibility, we took advantage of the unique VSG repertoires in the EATRO1125 and Lister427 parasite strains^2,23^. While AnTat1.1 is not endogenously present in Lister427 parasites, there are five VSGs nearly identical to the AnTat1.1 family member VSG-2986 (99.5%, 98%, 97.5%, 95.5%, and 92.2% nucleotide sequence identity) (3A, Supplemental Figure 4). These VSGs, if mosaic recombination were templated, could serve as donor VSGs for the formation of AnTat1.1-derived mosaic VSGs. We thus engineered dox-inducible Cas9-expressing Lister427 parasites and replaced VSG-2 with AnTat1.1 at the active expression site, BES1. We then induced breaks in AnTat1.1 as before and detected hundreds of recombination events in two independent clones (L1-A1 1 = 398, L1-A2 = 833; 3C). All mosaic recombination events detected utilized donor VSGs found exclusively in the Lister427 genome (3B), indicating that the repair events that generate mosaic VSGs require a template.

**Figure 3.**
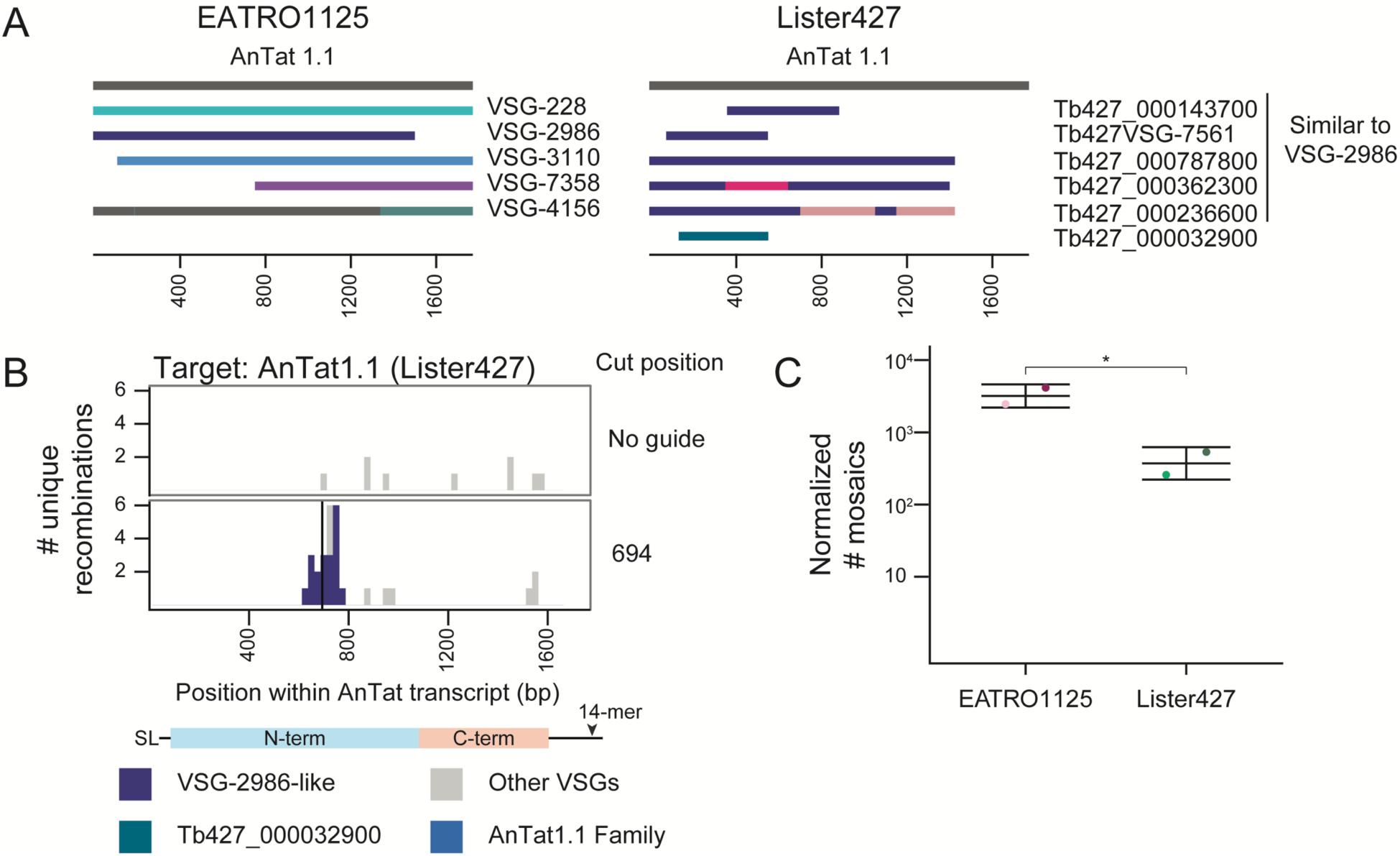
Nucleotide changes observed in mosaic VSGs arise from templated insertions. A) Schematics of the EATRO1125 AnTat1.1 VSG family and the VSGs similar to AnTat1.1 found within Lister427. Each is aligned to the AnTat1.1 transcript. For regions that are identical between two donor VSGs, regions are shown in the same color. B) A histogram of unique recombination events identified within AnTat1.1-expressing Lister427 parasites after a cut at position 694 along the AnTat1.1 transcript. The midpoint of the perfect homology between AnTat1.1 and the donor VSG at the recombination site is plotted. All AnTat1.1 family members except VSG-2986 were included as potential identifiable donor VSGs (all Lister427 VSGs + VSG-228, VSG-3110, VSG-7358, and VSG-4156). C) Quantification of mosaic recombination events induced by Cas9 at position 694. EATRO data is from 2A and 2B. The number of recombination events detected within 250bp up or downstream of the cut site was normalized to the number of total unanchored reads aligning within that region compared to the unanchored read count from the region with the smallest coverage from the EATRO cut sites to control for sequencing depth. Mean ± s.d. SL = 5′ splice leader sequence, 14-mer = 3′ sequence conserved in all VSG transcripts

### Donor VSGs can be utilized for mosaic recombination even within non-VSG loci

We wondered whether the location of a VSG within the genome could influence its utilization as a donor for mosaic recombination. To evaluate the role genomic context plays in the selection of donor VSGs, we again took advantage of the unique repertoires of the EATRO1125 and Lister427 strains^2^, as well as the high quality, phased genome assembly of the Lister427 parasite strain^23^. Using the same AnTat1.1-expressing Lister427 Cas9 parasites, we inserted an EATRO1125 VSG, VSG-228, which is commonly used as a donor in AnTat1.1 mosaics and is absent from Lister427 parasites, into three genomic locations: the minichromosomes, where VSGs are typically found, and the rDNA spacer and the tubulin array, where VSGs are not typically stored (Supplemental Figure 4). We then induced breaks at position 694 in AnTat1.1 and analyzed donor selection by VSG-AMP-seq (Figure 4).

**Figure 4.**
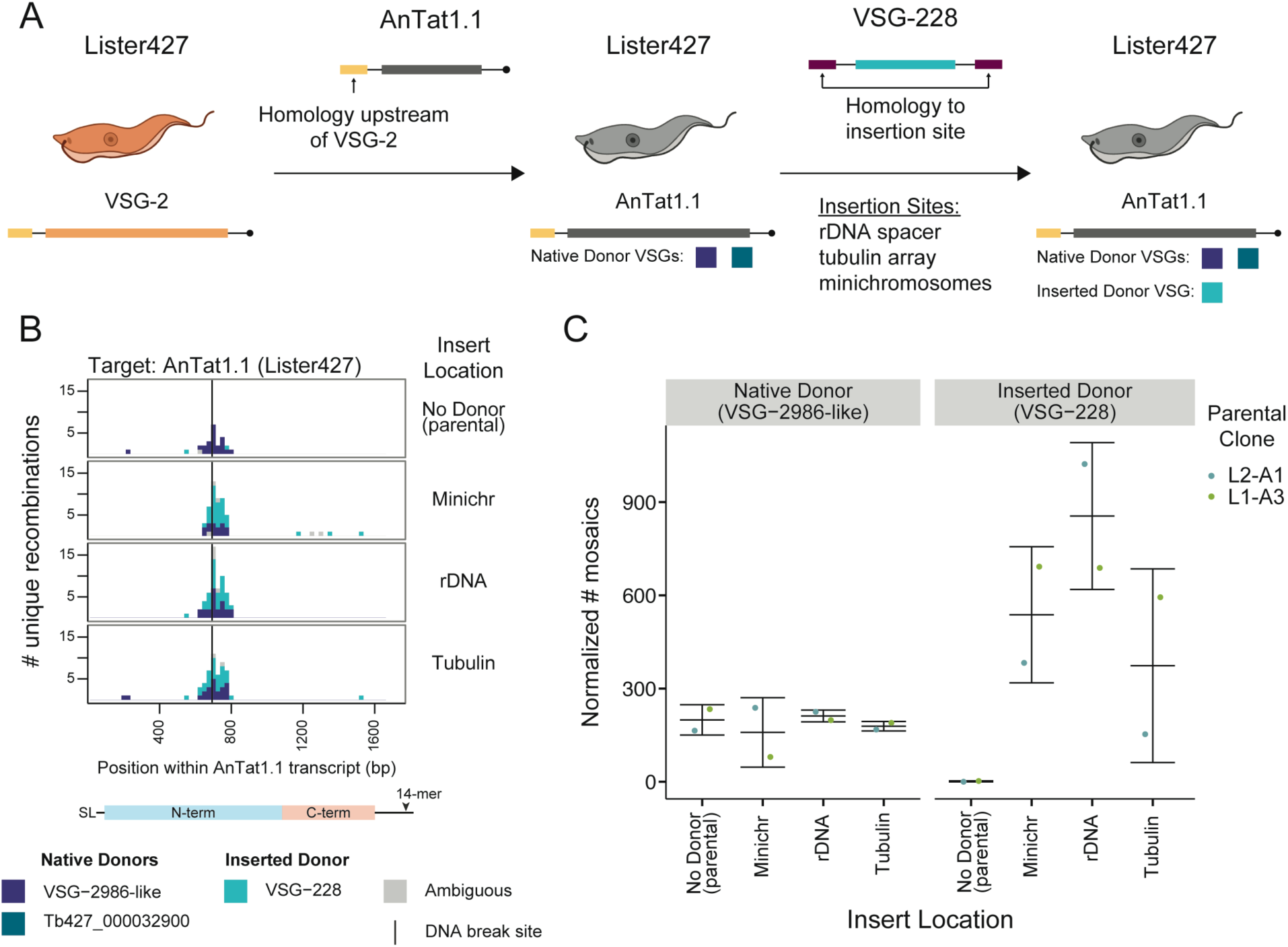
Donor VSGs are accessible throughout the genome. A) A schematic of the generation of Lister427 parasites expressing AnTat1.1 and harboring a silent copy of VSG-228. Available donor VSGs for the actively expressed VSG in each cell line are depicted below the parasites. B) A histogram of unique recombination events after a cut at position 694 in AnTat1.1 in Lister427 parasites expressing AnTat1.1 with VSG-228 inserted into the genome. The midpoint of the perfect homology between AnTat1.1 and the donor VSG at the recombination site is plotted. EATRO1125 AnTat1.1 family members were tested as potential identifiable donor VSGs (VSG-2986, VSG-228, VSG-3110, VSG-7358, and VSG-4156). C) Quantification of mosaic recombination utilizing VSG-2986-like or VSG-228 as a donor sequence at position 694 following induction of Cas9. The number of recombination events detected within 250bp up or downstream of the cut site was normalized to the number of total unanchored reads aligning within that region compared to the unanchored read count from the region with the smallest coverage. Each parental clone was independently generated. Mean ± s.d. SL = 5′ splice leader, 14-mer = 3′ sequence conserved in all VSG transcripts *T. brucei* cartoons created with Biorender.com

The endogenous Lister427 donors were selected at the same frequency in all samples, suggesting that the presence of an additional donor did not interrupt the formation of these mosaic VSGs. Surprisingly, the inserted donor, VSG-228, was utilized regardless of its location. Notably, only the coding sequence of VSG-228 was inserted into the Lister427 background. This indicates that neither the upstream 70-bp repeats typically found interspersed within the VSG archive^24–27^ or the 5’- and 3’-UTRs of the VSG are required for locating donor templates. Taken together, these data suggest that parasites possess a sophisticated homology search mechanism that allows donor VSGs to be utilized regardless of genomic location.

### Mosaic VSGs generated *in vitro* are identical to a subset of mosaic VSGs formed *in vivo*

To investigate whether the mosaic VSGs we detected after break induction *in vitro* reflected the events that occur naturally *in vivo*, we performed VSG-AMP-seq on parasites isolated from wildtype mouse infections, on day 15 post-infection, when AnTat1.1 has been mostly eliminated from the blood (Supplemental Figure 5B & 5C). We observed a strong C-terminal bias to all recombination events (316 recombination events across 7 mice). Very few sequences aligned to AnTat1.1 within the N-terminus, suggesting that most of these mosaic VSGs were complete replacements of the VSG N-terminal domain (5A; Supplemental 5A). These donor VSGs shared significantly less homology with AnTat1.1 than the AnTat1.1 family members, usually only sharing spans of 100bps or less of imperfect homology within the C-terminal region of the VSGs. Nevertheless, there was still a short (average = 12 bp) span of perfect identity between AnTat1.1 and the donor at the recombination site (5E). Since VSGs expressed by parasites at the second peak of parasitemia are antigenically distinct from AnTat1.1, we reasoned these N-terminal replacement VSGs ensured complete immune evasion since only a small portion of AnTat1.1 was retained.

**Figure 5.**
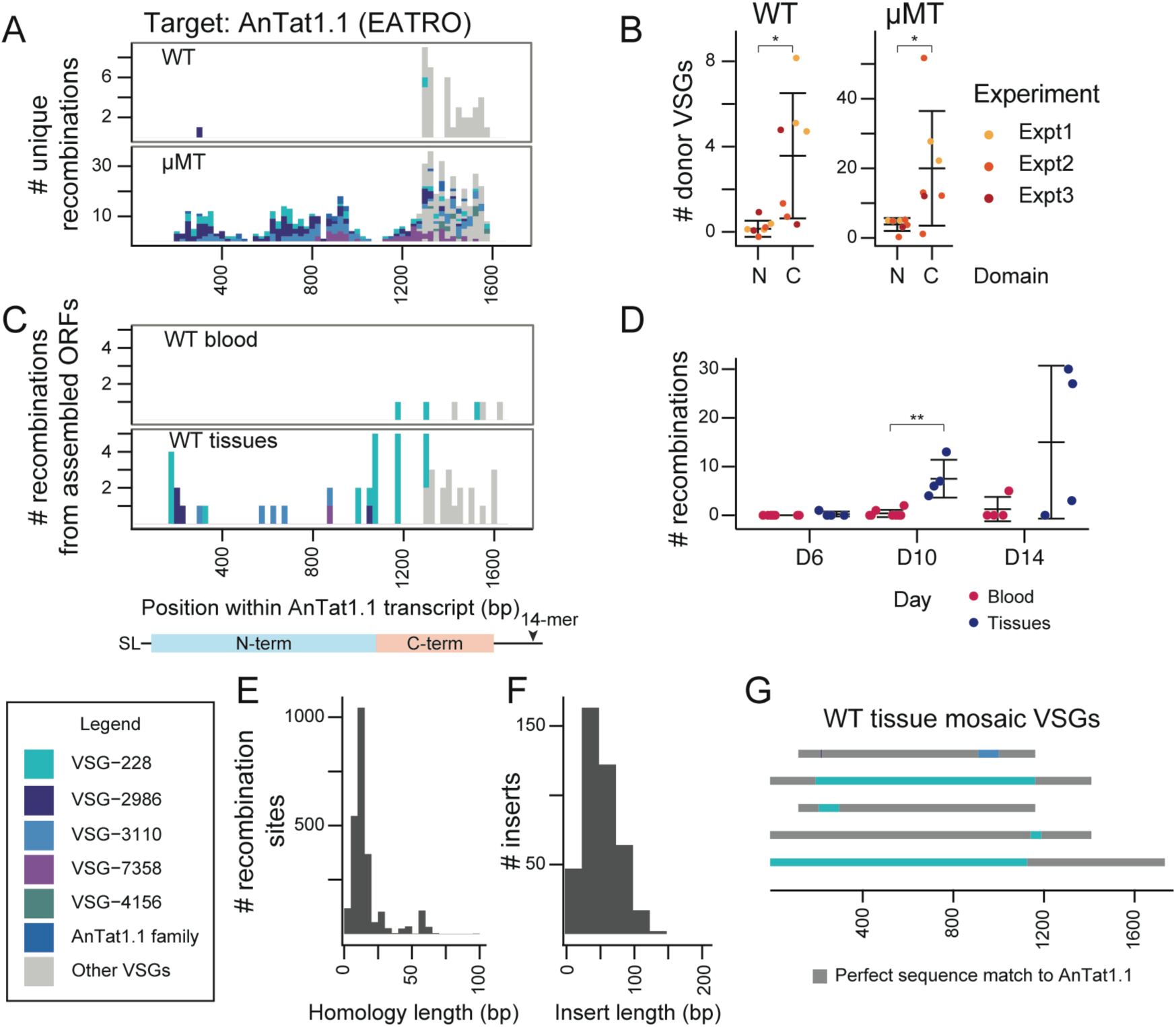
AnTat1.1 mosaic VSGs form *in vivo* and are most prevalent in extravascular spaces. A) A histogram of unique recombination events identified in all mice in wildtype or µMT mice from Day 15 post-infection. Recombination events found in multiple mice are represented once. (WT n=7; µMT n=7, from 3 independent experiments) The midpoint of the perfect homology between AnTat1.1 and the donor VSG at the recombination site is plotted. If a mosaic sequence matched >1 potential donor VSG, the average recombination position was plotted. B) Quantification of the number of donor VSGs utilized in mosaic recombination events within the N and C terminal domain of AnTat1.1 in wildtype and µMT mice from A). Statistical significance determined by a pairwise Wilcoxon test (*p<0.05) Mean ± s.d. C) A histogram of the mosaic recombination events identified from VSG ORFs assembled in Beaver et al.^4^ All recombination events identified are shown at all time points sampled during infection: D6, D10, and D14. If a mosaic recombination event was identified in more than one tissue within the same mouse, it was counted once (n=12, 4 mice per tissue time point). D) Quantification of the mosaic recombination events detected within wildtype mouse blood or tissue. Statistical significance determined by pairwise Wilcoxon within each timepoint. (**p<0.01) Mean ± s.d. E) A histogram of the length of the shared identity between AnTat1.1 and the donor VSG at the recombination sites. F) A histogram of donor VSG insertion lengths identified within individual reads. The insert length only includes newly inserted sequence and does not include recombination sites. G) Representative schematics of mosaic VSGs from assembled ORFs in Beaver et al.^4^ SL = 5′ splice leader sequence, 14-mer = 3′ sequence conserved in all VSG transcripts

We wondered whether the restriction of events to the VSG C-terminus reflected a mechanistic bias towards recombination within the VSG C-terminal domain or the effect of host antibody selection, with only the most immune evasive recombination events surviving. To investigate these possibilities, we repeated the experiment in µMT mice^28^, which do not have B-cells and therefore do not generate antibodies. We again analyzed day 15 post-infection, though AnTat1.1 is never cleared from these infections and parasitemia remains high throughout (Supplemental 5B & 5C). In µMT mice, mosaic recombination events span the full length of the VSG, and many are identical to mosaic VSGs observed *in vitro* (5A; Supplemental Figure 5A) (2144 recombination events across 7 mice). *In vivo,* we observe short insertions (average = 41 bp) using donors homologous to AnTat1.1 flanked by short (average = 13 bp, median = 9 bp) regions of perfect identity, matching the patterns we observed *in vitro* (5E & 5F). Interestingly, as in the wildtype infections, there are an additional subset of recombination events within the C-terminus which are largely absent *in vitro* and use a much more diverse set of donors (5B). These data demonstrate that our *in vitro* system recapitulates a subset of *in vivo* recombination events, but there may be other pathways facilitating recombination *in vivo* in addition to those that can be triggered by a single, blunt DNA double-strand break.

### Mosaic VSGs form in extravascular spaces during infection

While AnTat1.1 is eliminated from the blood in wildtype mice, it persists within tissues until at least day 14^4^. Given that parasite clearance is delayed in extravascular spaces, we wondered if mosaic formation may occur more readily in this parasite niche. To investigate this, we analyzed all assembled VSGs from a previous study of extravascular parasite populations^4^. Again, we found AnTat1.1 mosaics present in tissues identical to those observed in both µMT mice and *in vitro* (5C & 5G). Moreover, we observe that mosaic derivates of AnTat1.1 increase over time within tissue spaces during infection (5D). Together, these data suggest that mosaic VSGs form preferentially within extravascular spaces, possibly due to the slower VSG-specific parasite clearance in these spaces.

### Small changes in the VSG sequence provide substantial antibody evasion

Many AnTat1.1-derived mosaic VSGs differ from their original sequence by only a few amino acids. To determine how these changes impacted the antigenic character of the mosaic VSGs, we analyzed live parasites expressing AnTat1.1 mosaic derivatives by flow cytometry using a potent rabbit anti-AnTat1.1 polyclonal antibody raised against purified AnTat1.1 protein^29^ (6A). Many of the mutations block about 50% of antibody binding. Interestingly, three VSG-2986 mosaic clones were isolated with increasing insertion length (6B). The smallest insertion (Mosaic 2: 18bps, resulting in 3 a.a. changes), appears to account for most of the antigenic change, with only small decreases in binding associated with each additional length of insertion. We modeled the structure of these mosaics using ColabFold^30^ and their general structures matched AnTat1.1 (6C), except at the disordered top of the N-terminal lobe. A careful examination of the locations of the mutations on AnTat1.1 show that mutations from Mosaic 2 which have the largest effect on antibody binding are at the apex of the structure while other alterations can be found on the side of the monomer (6D). These results demonstrate that, although the insertions characteristic of mosaic recombination are quite short, even these small changes to the VSG can confer large consequences for host antibody binding depending on their position.

**Figure 6.**
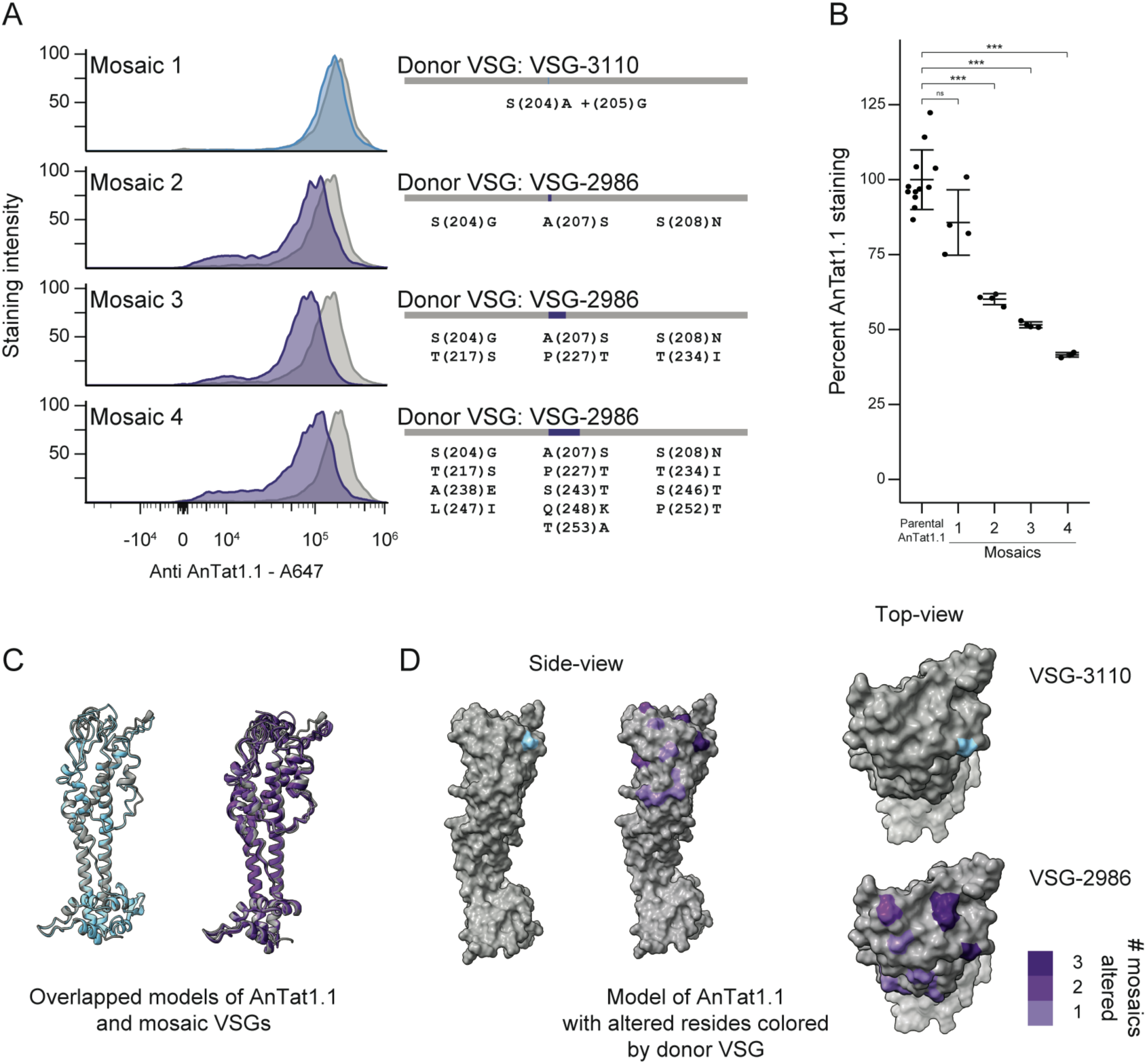
Mutations in sequences encoding the top of the VSG alter antibody binding. A) A histogram showing the binding of anti-AnTat1.1 antibody, as measured by anti-rabbit IgG Alexa Fluor 647 staining intensity, for parental controls (gray) and mosaic clones colored by donor VSG. (n= 4, from 3 independent parental lineages derived from 2 Cas9 clones) Schematics showing the mosaic VSGs are to the right. Donor VSG and amino acid substitutions are specified. B) Quantification of staining intensity changes for individual clones, based on median staining intensity. The median Alexa Fluor 647 intensities were normalized to the average of the parental clone. Statistical significance was determined based on a one-way ANOVA with a post-hoc Tukey HSD. (***p<0.001) Mean ± s.d. C) Overlapping ribbon structures of AnTat1.1 and Mosaic 1 in blue and AnTat1.1 and Mosaic 2, Mosaic 3, and Mosaic 4 in purple as predicted by ColabFold. D) A space filling model of AnTat1.1 highlighting the changed residues within the monomer. VSG-3110 in blue and VSG-2986 in purple.

## Discussion

Chronic *T. brucei* infection relies on the generation of new VSGs, with novel VSGs dominating late stages of infection^3,5,6^. However, the new VSGs generated during infection are complex, possibly arising from several intermediate events, and selected within a host environment with immunity against numerous previously expressed VSGs. The inherent complexity of chronic infection thus obscures the underlying biological principles driving VSG diversification. Here, we demonstrate that VSG diversification can be induced *in vitro* using Cas9-mediated double-strand DNA breaks within the VSG coding sequence, reproducibly generating mosaic VSGs that faithfully recapitulate mosaics formed naturally *in vivo*. By selecting just one VSG and looking at thousands of recombination outcomes, we have defined patterns characteristic of mosaic recombination. Mosaic VSGs typically form through short, templated insertions, and homology drives this process, restricting donor VSGs within the N-terminus to those from a set of closely related family members. Finally, we demonstrate that mosaic VSGs provide substantial immune evasion, particularly when these changes occur at the top of the VSG N-terminus, which may reflect a hypervariable region within the VSG.

We have shown that DNA breaks, previously shown to drive VSG switching^19,20^, can also result in VSG diversification through mosaic VSG formation. While the exact DNA repair mechanism that generates mosaics is unclear, the patterns we observe— short templated insertions that rely on short stretches of sequence homology—suggest this is not the canonical homologous recombination that mediates many VSG switch events. Homologous recombination relies on RAD51, an enzyme which binds free DNA ends at double strand breaks and stabilizes insertion into an intact DNA duplex, ultimately facilitating recombination. RAD51-dependent reactions in *T. brucei* require stretches of perfect homology of ∼100 bp^31^, and mismatches within the homology dramatically ablate recombination^31^. No recombination events detected here fit these restrictions; the maximum shared homology without a mismatch is 58 bp. In line with this, there is evidence for a RAD51-independent pathway capable of driving antigenic variation in *T. brucei*^32^. A set of *in vitro* studies in *T. brucei*^31,33^ suggested that the RAD51-independent recombination mechanism utilizes shorter homology sequences^31,33^ and more effectively tolerates mismatches between the target and donor sequences^31^. These patterns are more consistent with the recombination events we observe here, suggesting that mosaic formation may be driven by a RAD51-independent mechanism. While DNA repair utilizing short homology has been described in *T. brucei*^33–38^, the molecular details of the mechanism remain unknown. It will be critical to follow up on this study, given the tools we have now developed, to determine the precise mechanisms driving mosaic formation.

One surprising aspect of mosaic formation is the remarkably sensitive homology search mechanism capable of locating just 1600 bp of donor VSG sequence from genomic locations where VSGs are not typically stored. This was particularly unexpected given the structured organization of the VSG repertoire, with silent VSG genes primarily restricted to the subtelomeres^24–27^ and minichromosomes^39,40^ among interspersed fragments of 70 bp repeats. These conserved repeats have been thought to be critical mediators of the majority of VSG switch events by acting as upstream homology to VSGs within the silent subtelomeric and minichromosomal repertoires^41^. Ultimately, removing 70 bp repeat elements did not interrupt the homology search mechanism used for mosaic formation, suggesting the repeats are not required to identify a mosaic VSG donor template. This further reinforces our hypothesis that the repair mechanism driving mosaic formation is distinct from that driving VSG switching. With a search mechanism so sensitive, we suspect even small, extremely degenerate VSG fragments may be used by the parasite for DNA repair, resurrecting ancient VSG sequences for use in novel antigen formation.

While our data suggest most VSGs in the archive are capable of diversification through the mechanism we have described, this is not the case for all VSGs. Indeed, we find that mosaic recombination occurs only if a homologous donor VSG is available. Most studies of antigenic variation in *T. brucei* have focused on VSG-2, a unique VSG that does not readily diversify; this may explain, at least in part, why mosaic recombination has been observed so rarely *in vitro* until now. The proportion of VSGs within families is surprisingly high in both the EATRO1125 and Lister427 strains. Perhaps the VSG repertoire has evolved to provide a balance between diversification events, which allow significant expansion of the antigenic repertoire but may not always provide full immune evasion, and switching, which is more likely to completely escape pre-existing immunity.

While the members of the AnTat1.1 VSG family provide numerous possibilities for recombination and the generation of new VSGs, we found that the center portion of the gene appears to be especially successful at generating recombinants. We considered that this might be attributable to available donor sequences or guide efficiency, but there was nothing apparently unique about this site compared to other break locations. Protein modeling of AnTat1.1, however, revealed that this region encodes the top of the N-terminal lobe of the VSG, a site directly exposed to host antibody. For AnTat1.1 and many similar VSGs^2^ (A2-type), this is an unstructured region^42^, and indeed the apex of most crystalized VSGs appears unstructured despite a surprising variety of underlying structures^42^. We propose that this disordered region may have evolved to be more tolerant to amino acid changes, allowing diversification within the part of the protein most likely to facilitate immune evasion. While many recombination events may occur after a DNA break, we hypothesize that only a few, particularly those within disordered regions, maintain VSG structural integrity. Since expression of the VSG is essential for cell cycle progression^43^, it is possible that many newly formed mosaic VSGs lead to cell death for the parasites, allowing only a subset of the recombination events to survive and resulting in an apparent increase in recombination within the regions of the VSG where mutation is most tolerated.

Indeed, even very small changes within the top of the VSG N-terminal lobe can confer substantial immune evasion. Just three amino acid substitutions, from an 18 bp templated replacement, block approximately 50% antibody binding from a potent polyclonal antiserum. However, a 50% drop in binding may not be sufficient to fully evade preexisting anti-VSG antibody. Perhaps as a result, mosaic VSGs derived from AnTat1.1 in WT blood appear to be mostly N-terminal replacements where the AnTat1.1 N-terminal domain is completely replaced. A wider diversity of AnTat1.1-derived mosaics can be observed within the tissues, however, where AnTat1.1-expressing parasites linger due to differences in immune pressure^4^. In µMT mice, where AnTat1.1 expressing parasites can linger indefinitely, this effect is further amplified. We thus hypothesize that generating new VSGs is an iterative process, where a series of recombination events progressively shifts the character of a VSG until an immunologically distinct variant is formed. The relatively “protected” tissue spaces may serve as a site for this iterative process to occur. Another intriguing possibility is that partial immune evasion is advantageous for the parasite: a recent study suggested that exposure to sublethal antibody concentrations can trigger a VSG switch in *T. brucei*^44^. Perhaps parasites expressing these partially evasive variants are more prone to switching to a completely new VSG.

In addition to shedding light on the selective pressures imposed upon mosaic variants, our mouse data also demonstrate that our *in vitro* model of antigenic diversification recapitulates VSG diversification *in vivo*. The recombination events we detect in µMT mice represent the full breadth of possible recombination events which AnTat1.1 can form, and these are largely represented within our *in vitro* break-induced mosaic VSG populations. We observe short insertions flanked by short regions of identity in both contexts, but the dominance of the 694 break position we observe *in vitro* is not reflected in our *in vivo* analysis. This could be for a variety of reasons, including non-uniform break locations along the VSG *in vivo* or a variety of DNA break types driving a different pattern of outcomes. Intriguingly, in the absence of an exogenous DNA break, we do not observe any diversification *in vitro*, while diversification occurs continuously in the µMT context. This suggests that some aspect of the host environment other than antibody pressure induces diversification *in vivo.* It remains to be investigated whether an *in vivo* cue for diversification exists.

The diversification mechanisms we have described, where homology templated DNA repair drives antigen evolution, may also be at play in other gene families where diversity is critical. Many pathogens rely on diverse antigen genes to escape the host immune system, expressing one variable antigen from a silent repertoire stored within subtelomeric regions of the genome, which are inherently susceptible to DNA damage. Moreover, many eukaryotic pathogens that utilize antigenic variation (e.g. *Plasmodium*, *Giardia*, and *Trypanosoma*) have lost canonical non-homologous end-joining DNA repair^45^, suggesting an important role for alternative RAD51-independent repair mechanisms. Importantly, mosaic antigens have been observed in *Plasmodium falciparum* var genes^46–49^ and giardia variant-specific surface proteins (VSPs)^50,51^, some of which resemble the small insertion mosaics we have described here. Notably, the large subtelomeric multigene family of PIR genes in plasmodium^52,53^, which may drive antigenic variation in both *Plasmodium vivax* malaria and rodent malarias, lacks the repeat elements associated with most subtelomeric antigens^54^, a feature that is not required for this mechanism of DNA repair. Given these observations, we suspect the pathway described here may contribute to the evolution of antigen repertoires in many diverse organisms. Further, many alternative forms of DNA repair are also conserved in mammals^55^; it is thus plausible that large gene families like olfactory receptors^56,57^ and protocadherins^56^, which depend upon diversity and are known to swap sequences through gene conversion events^58–61^, may also be diversifying through similar mechanisms. A rigorous targeted approach, like we have used here, may be required to determine the role this mechanism plays in the diversification of other gene families.

Here we answer long-standing questions about how the *T. brucei* VSG repertoire evolved to generate new antigens critical for maintaining a chronic infection. Our high-throughput, sensitive technique has enabled the characterization of thousands of mosaic diversification events, orders of magnitude more than was previously feasible. This analysis revealed a pattern of short, homology-driven, and templated insertions around DNA break sites that can shift the antigenic character of the VSG. More broadly, this study provides an experimental framework for the hypothesis-driven exploration of antigen diversification, not only in *T. brucei* but also in other pathogenic microbes.

## Methods

### Lead contact

Further information and requests for resources and reagents should be directed to and will be fulfilled by the lead contact, Monica Mugnier.

### Materials availability

Plasmids associated with this study will be deposited to Addgene.

### Data and code availability

Raw sequencing reads from VSG-AMP-seq and VSG-seq was deposited to the NCBI Sequence Read Archive under project: PRJNA1140873

FASTQ files from assembled VSGs expressed by isolated parasite clones can be found at github.com/mugnierlab/Smith2024

Original code associated with this study can be found at github.com/mugnierlab/Smith2024

### Parasites

Pleiomorphic EATRO1125 AnTat1.1 90-13 *T. brucei* parasites were maintained in HMI-9 media with 10% heat inactivated FBS and 10% Serum Plus. Parasites were passaged when they reached approximately 5*10^5^ cells/mL. Monomorphic Single Marker Lister427 VSG221 TetR T7RNAP bloodstream form (NR42011; LOT: 61775530)^62^. *T. brucei* were maintained in HMI-9 up to 1*10^6^ parasites/mL. VSG221 has since been renamed to VSG-2.

### Plasmids

Plasmids were synthesized with Gibson Assembly. Gibson Assembly master mix was custom made^63^. Whole plasmid sequencing was performed by Plasmidsaurus using Oxford Nanopore Technology and their custom analysis and annotation pipelines.

pLEW100v5-BSD-FLAG-La-Cas9 was synthesized from pLEW100v5-BSD and pRPaCas9^64^. A *T. brucei* codon optimized FLAG tag was added to the N-terminus of Cas9. pLEW100V5-BSD was a gift from George Cross (Addgene plasmid #27658; http://n2t.net/addgene:27658; RRID:Addgene_27658). pRPaCa9 was a gift from David Horn (Addgene plasmid #111819; http://n2t.net/addgene:111819; RRID:Addgene_111819).

pHH-HYG-AnTat1.1 and pHH-HYG-VSG-228 were synthesized from pHH-HYG-VSG-3-S317A.^65^ This was a gift from Joey Verdi. AnTat1.1 sequence was obtained from AnTat1.1 specific cDNA from mouse infection D6 cloned into a pMiniT vector with the PCR Cloning Kit (NEB, E1202S). VSG-228 was partially amplified from VSG PCR (see below) derived from AnTat1.1 depleted D16 µMT parasites from blood. The partial fragment was amplified with an AnTat1.1 family specific forward primer (5’ - ACTACACCCACAACAAGCTCTA-3’) and a pan VSG reverse primer which binds to a conserved 14bp region of the 3’ UTR (5’-GATTTAGGTGACACTATAGTGTTAAAATATATC-3’) with AmpliTaq Gold (Applied Biosystems, 4398881) (anneal & extension 60C 1m, 35 cycles).The resulting PCR was cloned into the pMiniT backbone and the remainder of VSG-228 was *de novo* synthesized with Gibson Assembly.

The pT7sgRNA plasmids were obtained according to Rico et al.^64^ Briefly, sticky-end annealed sgRNAs were ligated into a BbsI-HF (NEB, R0539S) digestion of pT7sgRNA. Table 1 contains the sequences for the annealed guides. pT7sgRNA was a gift from David Horn. (Addgene plasmid #111820; http://n2t.net/addgene:111820; RRID:Addgene_111820)

**Table 1.**
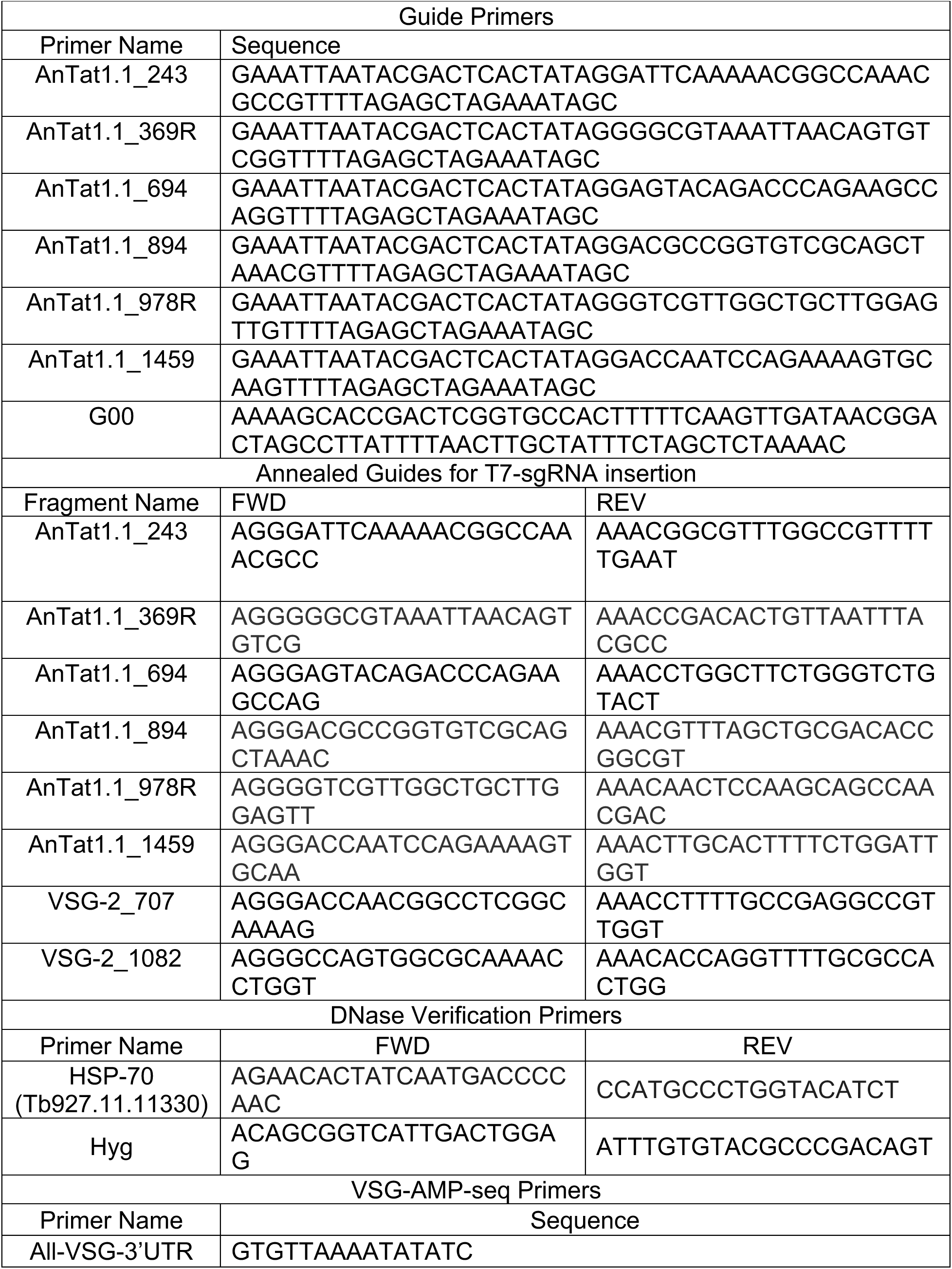
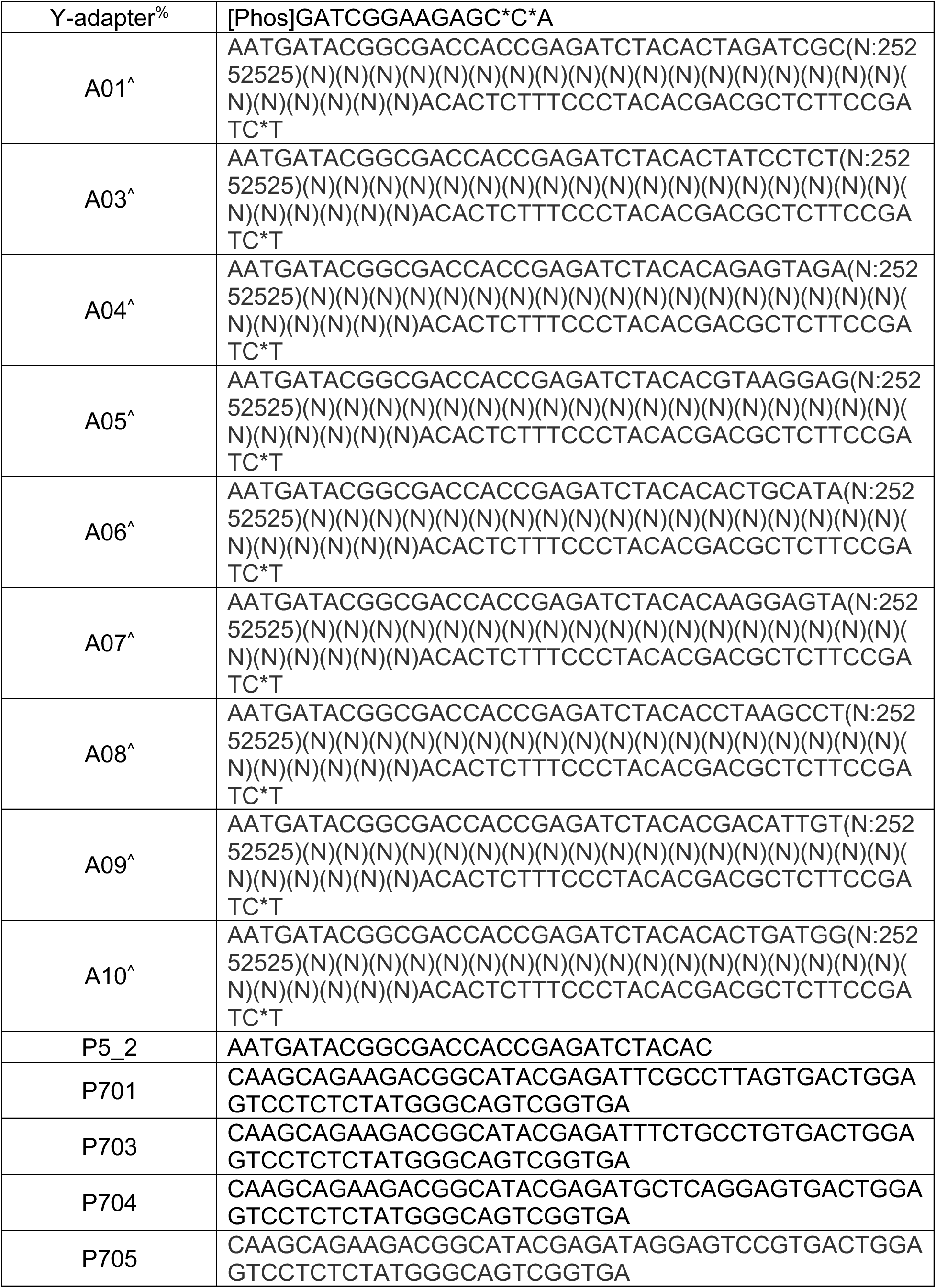
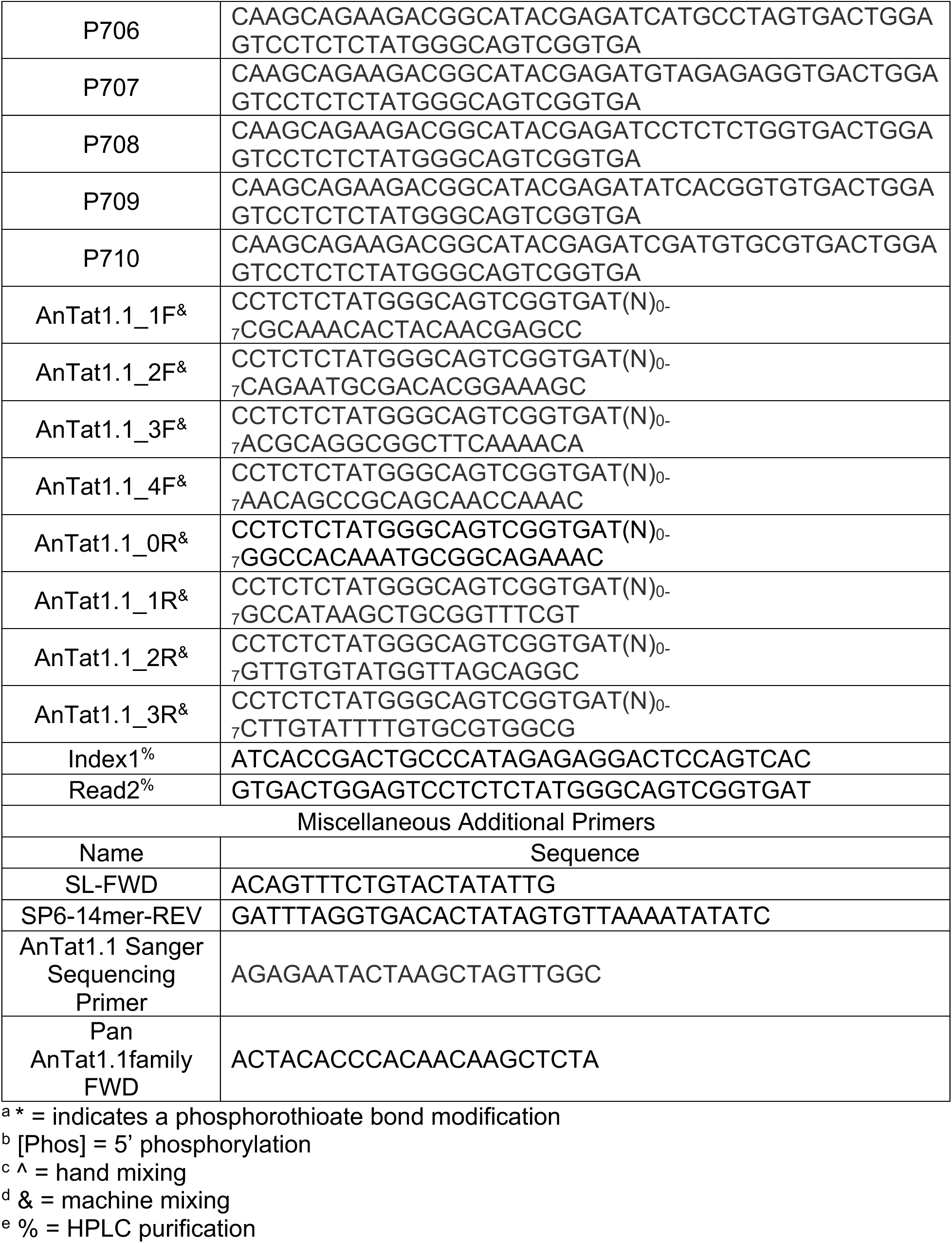
Guide and VSG-AMP-seq primers.

pHD309-PUR-VSG-228 was synthesized from pHD309-HYG-PUR. To obtain a plasmid for VSG-228 insertion, the sequence encoding the puromycin resistance gene was replaced with VSG-228. To convert the plasmid from hygromycin resistance to puromycin resistance, the sequence encoding the hygromycin resistance gene was replaced by the puromycin resistance gene. pHD309-HYG-PUR was a gift from George Cross. (Addgene plasmid #24014; http://n2t.net/addgene:24014; RRID:Addgene_24014)

pLEW100v5-PUR-VSG-228 was synthesized from pLEW100v5-BSD. The sequence encoding the blasticidin resistance gene was replaced with puromycin resistance. The sequence encoding the firefly luciferase was replaced with VSG-228. The rRNA promoter and the Tet operator sequences upstream of VSG-228 were removed.

pLEW100v5-PURO-177-VSG-228 was synthesized from pLEW100v5-177-HYG. The sequence encoding the hygromycin resistance gene was replaced with puromycin resistance. The sequence encoding firefly luciferase was replaced with VSG-228. The Tet operator sequences were removed. pLEW100v5-177-HYG was a gift from George Cross. (Addgene plasmid # 24013; http://n2t.net/addgene24013; RRID:Addgene_24013)

pLEW100v5-PURO-T7-sgRNA was synthesized from pLEW100v5-PUR-VSG-228 and pT7sgRNA. 5’ UTRs for Fructose 1,6-bisphosphate aldolase replaced the Actin 5’ UTR for puromycin expression. Actin and GPEET 5’UTRs were removed as they have BbsI restriction sites within them. A T7 promoter was added to drive PURO expression. The T7-sgRNA cassette from pT7sgRNA was inserted into the modified pLEW backbone. Target specific constructs were obtained as above for pT7-sgRNA.

### Transgenic parasites

To obtain all transgenic parasites, 5 million parasites were electroporated with 5-10ug of digested plasmid with an AMAXA Nucleofector II using X-001 in Human T-cell Nucleofector Solution (Lonza VPA-1002).

For tetracycline-inducible Cas9 parasites, EATRO1125 or Lister427 parasites were electroporated with pLEW100v5-BSD-FLAG-La-Cas9, digested with NotI-HF (NEB, R3189S). Parasites were immediately selected in 5ug/mL blasticidin (Thermo Scientific, R21001) and maintained in selection unless otherwise specified.

To obtain AnTat1.1 and VSG-228 expressing Lister427 parasites, Lister427 parasites were electroporated with pHH-HYG-VSG plasmids digested with BamHI-HF (NEB, R3136S). After 16-24 hours recovery, 25ug/mL of hygromycin (Fisher Scientific, J67371-XF) was added to the culture. After obtaining colonies 5-7 days later, selection was reduced to 5ug/mL and maintained unless otherwise specified.

Lister427 parasites with pLEW100v5-BSD-FLAG-La-Cas9 inserted were additionally electroporated with pHH-HYG-VSG plasmids expressing Antat1.1. Clones were obtained as above.

To obtain AnTat1.1 Lister427 parasites with silent VSG-228 donors inserted, these parasites were additionally electroporated with pHD309-PUR-VSG-228, pLEW100v5-PURO-VSG-228 or pLEW100v5-PURO-177-VSG-228 plasmids digested with NotI-HF. After 16-24 hours of recovery, 0.1ug/mL of puromycin (Millipore Sigma, P8833) was added to the culture. Parasites were maintained in puromycin, and 5ug/mL of blasticidin and hygromycin unless otherwise specified.

To obtain constitutive guide expressing parasite lines, inducible Cas9-expressing parasites (both Lister427 and EATRO1125 Cas9 strains) were electroporated with pT7sgRNA guide containing plasmids digested with NotI-HF^64^. After 16-24 hours recovery, single colonies of parasites were selected with 2ug/mL phleomycin (Sigma-Aldrich, SML3001). Parasites were maintained without phleomycin selection. These parasites were used to obtain parasite clones following DNA break induction.

To obtain constitutive guide expressing parasite lines with puromycin selection rather than phleomycin, EATRO1125 inducible Cas9-expressing parasites were electroporated with pLEW-T7-sgRNA digested with NotI-HF. After 16-24 hours of recovery single colonies were selected with 0.1ug/mL of puromycin. Cultures were maintained in 0.1ug/mL puromycin and 5ug/mL blasticidin unless otherwise specified. These parasites were used for western blot analysis and to obtain parasite clones following DNA break induction.

### T7-Guide synthesis & purification

DNA fragments containing a T7 promoter and guide RNA sequence were synthesized as described previously^66^. Briefly, T7-guide primers and a G00 primer were amplified with Phusion polymerase for 35 cycles (annealing temp 60C, extension 5s). PCR products from 12 identical PCRs were pooled and purified via ethanol precipitation.

### Cas9 transient electroporations

Approximately 24 hours prior to electroporation, parasites were seeded at a density of 83,000 parasites/ml and induced with 1ug/mL doxycycline (Millipore Sigma, D9891-1G). The total culture volume of was determined by the number of samples being electroporated; 12mLs were seeded per sample. Blasticidin selection (Cas9) and puromycin selection (silent VSG-228) were maintained, if applicable, and for Lister427 parasites expressing AnTat1.1, hygromycin (active VSG expression) was removed at this stage. Either 8mLs (Lister427) or 10mLs (EATRO1125) of the bulk parasite culture, approximately 5-10 million cells, were spun down for each sample. Media was then removed, and parasites were resuspended in 100uL Human T-cell Nucleofector Solution. Each sample was electroporated using the X-001 program on the AMAXA Nucleofector II with approximately 1-1.5ug of purified T7-guide in a volume less than 10uL or a sample without any DNA as a negative control. Parasites were moved into 5mLs HMI-9 in 6-well plates to recover for 30 minutes then moved into 20mLs total in flasks to recover overnight. About 24 hours after electroporation, parasites were counted and split. For EATRO1125, 12 million cells (or all cells if there were fewer than 12million) were seeded into 60mLs total with blasticidin. For Lister427, 2 million cells were seeded into 20mLs total with blasticidin. At the 48 hour time point, parasites were counted, collected, and stored in TRIzol (Invitrogen, 15596026) for subsequent RNA extraction.

### Isolation of mosaic expressing parasites

Multiple parasite clones expressing T7sgRNA with guides at cut sites 243 and 694 were derived from two distinct inducible Cas9-expressing parasite clones with constitutive guide expression driven by pT7-sgRNA. Multiple dilutions of parasites were plated in 96-well plates in 5ug/mL blasticidin and 1ug/mL doxycycline. Parasite clones were isolated from plates containing fewer than 30 individual surviving parasite clones. Most plates contained fewer than 10. Individual clones were visible after 7-14 days. Upon isolation of a parasite clone, all drugs were removed. A subset of parasites were analyzed by flow cytometry.

A single parasite clone expressing pLEW-T7-sgRNA with a guide at position 1459 was derived from an inducible Cas9-expressing parasite clone. Multiple dilutions of parasites were plated in 96-well plates in 5ug/mL blasticidin, 0.1ug/mL puromycin, and 1ug/mL doxycycline. Parasites were isolated from plates containing fewer than 30 surviving parasite clones. Individual clones were visible after 7-14 days. Upon isolation of a parasite clone, all drugs were removed.

VSG sequences for clones were determined from extracted RNA. cDNA was synthesized using the Superscript III Reverse Transcriptase (Invitrogen, 18080051) and a VSG-specific primer which binds to a conserved 14-bp sequence within the 3’ UTR. (5’-GTGTTAAAATATATC-3’). 2uL of RNase-treated cDNA was amplified for 35 cycles with VSG specific primers: a spliced-leader (5’-ACAGTTTCTGTACTATATTG-3’) and SP6-VSG 14-mer (5’-GATTTAGGTGACACTATAGTGTTAAAATATATC-3’) using Phusion polymerase (Thermo Fisher, F530L) (annealing temp 55C, extension 45s). PCR products were cleaned using the Monarch PCR & DNA cleanup kit (NEB, T1030L). VSG sequences were determined by amplicon nanopore sequencing performed by Plasmidsaurus using Oxford Nanopore Technology with their custom analysis and annotation where fragmented VSG amplicons were sequenced and assembled into full length sequences or targeted Sanger sequencing with the sequencing primer (5’-AGAGAATACTAAGCTAGTTGGC-3’) performed by Azenta Life Sciences.

### Mouse infections

All experiments involving mice were performed in accordance with the protocol approved by the Institutional Animal Care and Use Committee at Johns Hopkins University. 8-12 week old female C57BL/6 mice and µMT^-^ (B6.129S2-Ighm^tm1Cgn^/J)^28^ mice (Jackson Labs) were infected with ∼5 EATRO1125 parasites by intravenous injection in the tail vein. These parasites express either AnTat1.1 or VSG-421. Starting at D4 post-infection, parasitemia was monitored within mice every two days via tail bleed. Blood was harvested at D6 post-infection through a submandibular bleed. An additional gel pack and in-cage food pellets were provided to mice during recovery. At D15 post-infection, mice blood was harvested by cardiac puncture. Blood was stored in TRIzol LS (Invitrogen, 10296028) for RNA extraction.

### RNA preparation

Parasites were stored in TRIzol prior to RNA extraction. Blood with parasites was stored in TRIzol LS. RNA was extracted via phenol/chloroform extraction according to the manufacturers protocol. Purified RNA was DNase treated with Turbo DNase (Thermo Fisher, AM2239) and purified with 1.8X Mag-Bind TotalPure NGS Beads (Omega Bio-tek, M1378-01). Verification of effective DNase treatment was performed via PCR of hygromycin (EATRO1125 only) (Fwd: 5’-ACAGCGGTCATTGACTGGAG-3’; Rev: 5’-ATTTGTGTACGCCCGACAGT-3’, annealing temp 52C, extension 30s) or HSP70 (Lister427 & EATRO1125) (Fwd: 5’-AGAACACTATCAATGACCCCAAC-3’; Rev: 5’-CCATGCCCTGGTACATCT-3’, annealing temp 50C, extension 15s) genes for 30 cycles using OneTaq DNA Polymerase (NEB, M0480L).

### VSG-seq

VSG-seq was performed as previously described^3,4^. Briefly, cDNA was synthesized using the Superscript III Reverse Transcriptase and a VSG-specific primer which binds to a conserved 14-bp sequence within the 3’ UTR. (5’-GTGTTAAAATATATC-3’). cDNA was treated with Rnase A (Qiagen, 19101) and RNase H (Invitrogen, 18080051) for 30 minutes then purified with 1.8X Mag-Bind Total NGS Beads (Omega Bio-tek, M1378-01). Purified cDNA was amplified for 25 cycles with VSG specific primers: a spliced-leader (5’-ACAGTTTCTGTACTATATTG-3’) and SP6-VSG 14-mer (5’-GATTTAGGTGACACTATAGTGTTAAAATATATC-3’) using Phusion polymerase (Thermo Scientific, F530L) (annealing temp 55C, extension 45s). This PCR product was prepared for sequencing following a 0.6X bead clean up using the Nextera XT DNA Sample Prep Kit (Illumina, FC-131-1096) according to the manufacturer’s instructions. Libraries were sequenced with 100 bp single-end reads on a NovaSeq6000. Analysis was performed using the VSGSeqPipeline found at github.com/mugnierlab.

### VSG-AMP-seq library preparation

VSG-AMP-seq was based upon AMP-seq^67^ and GUIDE-seq^68^. cDNA was synthesized from DNA-free RNA using Superscript III Reverse Transcriptase and a VSG-specific primer which binds to a conserved 14-bp sequence within the 3’ UTR. (5’-GTGTTAAAATATATC-3’). Second Strand synthesis was performed with NEBNext mRNA Second Strand Synthesis Module (E6111L). The resulting double stranded cDNA was purified with 1.8X Mag-Bind Total NGS Beads. cDNA was fragmented briefly with NEBNext dsDNA Fragmentase (M0348L) for ten minutes at 37C to obtain fragments of approximately 500bps. Fragments were purified with a double-sided Mag-bind bead cleanup. First, fragments were incubated with 0.5X beads, DNA bound to the beads were discarded and an equal volume of beads as before, then a 1X PEG concentration, was added to the fragments and the cleanup proceeded as normal. Fragments were end-repaired with Enzymatics low concentration end repair mix (Y9140-LC-L) and A-tailed with Recombinant Taq (Life Technologies, 100021276). Y-Adapters were pre-annealed by incubating a MiSeq Common Adapter with Adapters (A01-A10) containing 25 bp Unique Molecular Indexes (UMIs) and a barcode at 10uM at 95C for 1 sec, 60C for 1s, and slowly cooled to 4C. Adapters were ligated to the A-tailed fragments with T4 DNA Ligase (Enzymatics L6030-LC-L). The resulting fragments were cleaned up with 0.8X Mag-Bind beads. Two target specific PCRs were performed on the fragments with pools of target specific primers, one in the forward and one in the reverse direction. All primers are listed in Table 1. Target primers have spacers of varying lengths to generate sequence diversity without the need for PhiX. Target specific primers are paired with P5_2 and a sample specific P primer (P701-P710) which contains a second barcode. Fragments are amplified with Platinum Taq (Life Technologies, 10966018) using the following program: 95C 5 mins, 35 cycles of 95C for 30s, 55C for 30s, and 72C for 30s, and 72 for 5 mins. The resulting products were cleaned up with 0.6X Mag-bind beads. Libraries were quantified using the Qubit dsDNA HS Kit (Life Technologies, Q32854) and run on a 1% agarose gel to determine average length. Libraries were sequenced on a MiSeq or for deep sequencing of µMT samples on a NovaSeq6000 with custom index1 and read2 primers using the following cycle conditions: “151|8|33|131” with the paired-end Nextera sequencing protocol.

### VSG-AMP-seq analysis

Index sequences were added to the names of sequencing reads. FASTQ files were demultiplexed by target specific primer using cutadapt v=3.5^1^ searching for multiple target-specific primers at the 5’ end of read2 in paired-end mode using flags -- action=retain --overlap 10. Then, reads were quality trimmed with trim_galore v0.6.4_dev (github.com/FelixKrueger/TrimGalore) where adapter sequences, if present, were removed with flags --dont_gzip --paired --trim1. Spacers were removed from read1 if present using cutadapt by searching for the target-specific primer sequence at the 3’ end of the read with flags --action=retain --overlap 10. Dual-indexed barcodes were used to demultiplex reads. A custom function in the pipeline (barcode_errors()) was used to the identify number of mismatches permitted so all barcodes present could be unambiguously identified.

To consolidate reads, UMIs were extracted from large FASTQ files, split into smaller files and grouped by 100% identity using cd-hit-est v4.8.1^70,71^ with flags -c 1.0 -n 8 -M 16000 -d 0. UMIs were then grouped by 92% (no more than 2 mismatches per UMI) using flags -c 0.92 -n 8 -M 16000 -d 0. Reads were sorted into consensus groups based on UMIs and a consensus sequence was formed as the most popular base at each position of the read with a threshold quality score of at least 2. If there was a tie, an N was used. Clusters with fewer than 3 reads were removed. Consolidation was only used for samples isolated from mouse infections.

The full sequence of the AnTat1.1 transcript was determined by Plasmidsaurus using Oxford Nanopore Technology with their custom analysis and annotation. However, in our initial analyses of control samples, we determined that many reads slightly differed near the splice leader, suggesting that transcripts might be alternatively spliced or included extra sequence appended at some point during the library preparation process, specifically adjacent to splice leader. We defined the AnTat1.1 reference as the full-length variant determined by Plasmidsaurus sequencing, with the longest splice variant observed in control samples (no guide, electroporated) appended to the 5’-end. When reads did not align to this reference, they were then compared against the ten most commonly observed AnTat1.1 isoforms to account for all possible variations in the AnTat1.1 transcript. These sequences, including the full length AnTat1.1 reference, can be found in global_target.py. The positions were 0 indexed. Read2, the anchored read, was aligned to AnTat1.1. Read pairs were removed if: the first 4/10 bases after the primer did not align to AnTat1.1, an alternative VSG which contained the primer sequence was amplified, the wrong position of AnTat1.1 was amplified by the anchor, if the reads were too short (<15 bps), contained too many Ns (>5), if there was an inversion event or a duplication, or if the transcript was alternatively spliced. Read1, the unanchored read, was also aligned to AnTat1.1 with a mismatch of up to 1bp. If no alignment could be found, read1 was trimmed to remove AnTat1.1 like sequences from the 5’ and 3’ ends, allowing up to 1 mismatch on each end to be removed, leaving a fragment which can be used to search the VSGnome and identify potential donor VSGs. A consensus sequence was generated from the reads, if they overlapped, using their alignment positions within AnTa1.1. AnTat1.1, the consensus read, and putative donor VSG were aligned, and recombination sites were identified. If no consensus could be generated from the two reads, read1 was used. Ambiguous recombination sites where the mosaic portion of a read matched more than one donor were represented by the average position of the potential recombination sites and identified as ambiguous. Mosaic reads which did not overlap to generate a consensus and did not contain an identifiable recombination site in read1 were excluded from analysis.

### Identification of mosaic VSG from ORFs in Beaver et al.^4^

ORFs from Beaver et al.^4^ were tiled into 20 bp k-mers overlapping by 1bp each. All tiles were aligned to AnTat1.1 using bowtie^72^ with the following flags: -v 2. ORFs where 15 or more tiles successfully aligned to the AnTat1.1 sequence were identified as possible mosaics (≥ 35 bp match, potentially discontinuous). AnTat1.1 sequence containing ORFs were assessed for sequence matching common donor VSGs: VSG-228, VSG-2986, VSG-3110, and VSG-7358. If a best match could not be determined or if recombination did not occur with one of the typical donor VSGs, VSG sequences were analyzed by hand to identify remaining mosaic recombination events.

### VSG clustering and family identification

VSGs were identified from annotated genes and pseudogenes from the TriTrypDB-66 Lister427 2018 genome GFF^23^. Those identified with an attribute including “VSG” or “variant” but not “invariant”, “Invariant”, “histone”, “Histone”, “RING”, “ubiquitin”, ”exclusion”, “Exclusion”, “nonvariant”, or “Nonvariant”. Many known VSGs were only annotated as an unknown protein product. To identify additional VSG-encoding genes, we isolated all unknown gene products with attributes including “hypothetical protein”, “pseudogene” or “unknown”, but not including “VSG”, “variant” or any of the additionally excluded words above. VSGs were identified among these unknown genes via BLAST against all known VSG-encoding sequences (Those from Cross et al.^2^, the EATRO1125 VSGs with flanking sequences from George Cross, Beaver et al.^4^, and TriTrypDB-66^73^ TRU927, Lister427, and EATRO1125 genomes.) Genes were considered a VSG if the blast hit extended over 80% of the unknown gene and had a bitscore of 500 or greater. We identified an additional 2545 VSGs.

All Lister427 identified VSGs from the Lister427 2018 genome and the FASTAS from Cross et al.^2^ (vsgs_tb427_all_atleast150aas_cds.txt, vsgs_tb427_nodups_atleast250aas_cds.txt, vsgs_tb427_nodups_atleast250aas_cdsplusflanks.txt) were combined and duplicate VSGs were removed with cd-hit-est with the following flags: -c 1.0 -n 8 -M 16000 -d 0. VSGs with identical sequences, but distinct positions within the 2018 genome were added back for a total of 8564 VSGs. In the Lister427 2018 genome, we identified 5789 VSG-encoding genes total and included 300bps flanking sequences up and downstream of the coding sequence for further analysis. These are genes visualized in Supplemental Figure 4.

EATRO1125 VSGs were obtained from TrypsRU (George Cross) and included 200 bp flanking sequences. (vsgs_tb1125_all_atleast150aas_cdsplusflanks.txt) Duplicate genes were removed with cd-hit-est with the following flags: -c 1.0 -n 8 -M 16000 -d 0.

To define VSG families, network analysis was performed using the curated VSG reference FASTA files described above. Each genomic repertoire was subjected to an all-versus-all BLASTn run under the default parameters to generate pairwise tables which contained the query-subject pair, query sequence length, alignment length, E-value and percent identity. Networks were generated with the igraph R package^74,75^ using undirected and unweighted clustering of nodes after applying link cutoffs based on E-value < 1e-20 and alignment coverage of the query sequence > 80%. The leading eigenvector clustering method (function: cluster_leading_eigen()) was used to detect communities and assign nodes based on their connectivity.

To supplement the BLAST network graph approach, the greedy clustering UCLUST algorithm^76^ was used to assess clusters of VSGs. VSGs were clustered using usearch with the following flags: -id .75 -strand both -sizeout -sort length -maxhits 2. For each genome, identified VSGs were sorted by sequence length then clustered at a global identity threshold of 75%

### Western blotting

EATRO1125 parasites with pLEW-FLAG-La-Cas9 and pLEW-T7sgRNA were grown for 24 hours in the presence of doxycycline or DMSO vehicle control. Post induction, 5 million parasites were harvested and washed with 25C PBS. Then, pelleted parasites were resuspended in 50uL RIPA buffer (50mM Tris, 150mM NaCl, 1% NP-40, 0.5% Sodium Deoxycholate, 0.1% SDS, pH = 7.4) + 2X Laemmli buffer. Lysed parasites were boiled at 95C for 5 minutes. 5uL of lysates were separated on a Tris-glycine polyacrylamide gel at 110V for 100 minutes in Tris/glycine running buffer (25mM Tris, 192mM glycine, 0.1% SDS, pH = 8.3). Proteins were then transferred onto a nitrocellulose membrane using transfer buffer (25mM Tris, 192mM glycine, 20% methanol) overnight at a 60mAmps per transfer box at 25C. For Cas9 (1:1000) and EF1α (1:1000), lysates were separated on 10% polyacrylamide gels and transferred onto 0.45µm membranes. For γ-H2A (1:200), lysates were separated on a 15% polyacrylamide gel and transferred onto a 0.2µm membrane. After transfer, blots were blocked with 0.5% BSA in TBS with 0.05% Triton X-100 (TBST, 20mM Tris, 150mM NaCl, pH = 7.6) for an hour at RT under constant agitation. Blots were incubated with primary antibody (see above dilutions) for 2 hours at 25C. After 5 TBST washes, blots were stained for 1 hour with goat anti mouse (1:5000, Cell Signaling, 7076S) or goat anti rabbit-HRP-conjugated secondary (1:5000, Cell Signaling, 7074S). After another 5 washes, blots were incubated with ECL (Cytiva, RPN2109) and film was exposed to blots in a dark room. Developed film was scanned and images were processed with FIJI^77^. Primary antibodies used were mouse anti FLAG (M2 clone) (Millipore Sigma, F3165-1MG), mouse anti EF1α (CBP-KK1 clone) (Millipore Sigma, 05-235), and rabbit anti γ-H2A was a kind gift from Galadriel Hovel-Miner based upon Glover and Horn^78^.

### Flow cytometry

In 96 well plates, 200,000 parasites were stained with 1:20,000 rabbit anti AnTat1.1 primary antibody^29^ (Jay Bangs) for ten minutes at 4C while shaking in PBS + 10mg/mL glucose. Parasites were washed once with 100uL PBS + glucose. Then, parasites were stained with Alexa Fluor 647-conjugated goat anti rabbit IgG (H+L), F(ab’)2 Fragment (Cell Signaling, 4414S) at 1:1000 at 4C while shaking in PBS + glucose. After washing again with 100uL PBS + glucose, parasites were resuspended in PBS + glucose + 1:20 Propidium Iodide (BD Biosciences, 556463) and analyzed on a Attune Nxt flow cytometer (Invitrogen). Data analysis was performed using FlowJo v10.

### Analysis and modeling of VSG N-terminal domains

Full length protein coding sequences from AnTat1.1 and its isolated mosaics were used for structural modeling. Only N-terminal domain sequences were used for protein structural prediction since this region of the VSG is the most well-defined experimentally. Signal peptides are cleaved from the mature VSG during processing, so we used SignalP 6.0^79^ to predict and remove the Sec/SPI sequence (--organism eukarya, --mode fast) resulting in a FASTA file of mature proteins. To determine the coordinate of the N-terminal domain, we used a python analysis pipeline available at (https://github.com/mugnierlab/find_VSG_Ndomains). The script identifies the boundaries of the VSG N-terminal domain using the HMMscan function under HMMer version 3.1b2^80^. Query sequences are searched against an HMM profile containing 735 known N-terminal domain sequences from Cross et al.^2^ and N-terminal domains defined by the largest envelope domain coordinate that meets E value threshold (1 x 10^-5^, -domE 0.00001). The processed FASTA file containing only mature VSG N-terminal domain sequences was used as input for structural prediction of monomers using LocalColabFold^30^ function colabfold_batch run using the following arguments: --amber, --templates, --num_recycle 3. The best ranked output model with the highest average predicted local distance test score (pLDDT), that is the highest confidence model, was used for downstream analyses. Model visualization and alignment were performed using with UCSF ChimeraX version 1.7.1^81^, developed by the Resource for Biocomputing, Visualization, and Informatics at the University of California, San Francisco, with support from National Institutes of Health R01-GM129325 and the Office of Cyber Infrastructure and Computational Biology, National Institute of Allergy and Infectious Diseases.

## Acknowledgements

We would like to thank Jay Bangs for generating and sending us primary rabbit anti-AnTat1.1 polyclonal antibody. Thanks to Galadriel Hovel-Miner for the primary rabbit anti-γ-H2A antibody. We appreciate members of the Mugnier lab for their thoughtful review of this manuscript, particularly Carolina Duque and Sneider Gutierrez. We also would like to thank Richard McCulloch for thoughtful feedback about this manuscript. We would like to thank David Mohr and the JHU GCRF Genomics Core for sequencing services. Thanks to Joey Verdi for pHH-HYG-VSG-3-S317A and advice about the generation of transgenic parasites. Data analysis was carried out at the Advanced Research Computing at Hopkins (ARCH) core facility (rockfish.jhu.edu), which is supported by the National Science Foundation (NSF) grant number OAC 1920103. This work was supported by the JHU Catalyst Award. JES, JMCH, AKB, JS, BZ, and MRM were supported by Office of the Director, NIH (DP5OD023065). JES and JS were additionally supported NIH training grant T32GM007 and JES was supported by the National Science Foundation (NSF) Graduate Research Fellowship under grant numbers 1232825 and 1746891. JMCH, AKB, JS and EHA were additionally supported by NIH T32AI007417.

## Author Contributions

Conceptualization, J.E.S. and M.R.M.; Methodology, J.E.S., J.M.C.H., A.K.B., A.M., S.D.G., B.Z., and E.H.A.; Software, J.E.S., A.M., J.S., and L.M.; Validation, J.E.S. and K.J.W.; Formal Analysis, J.E.S.; Investigation, J.E.S., K.J.W., E.M.K., J.S., A.K.B., A.M., J.Z., and D.N.M.; Resources, M.R.M.; Data Curation, J.E.S., A.M., and J.S.; Writing – Original Draft, J.E.S. and M.R.M.; Writing – Reviewing & Editing, J.E.S., A.K.B., and M.R.M.; Visualization, J.E.S. and J.S.; Supervision, J.E.S. and M.R.M.; Project Administration, J.E.S.; Funding Acquisition, J.E.S. and M.R.M. VSG-AMP-seq developed by J.E.S., S.D.G., and B.Z. EATRO1125 Cas9 experiments performed by J.E.S. Lister427 Cas9 and Lister427-Cas9-AnTat1.1 experiments performed by J.E.S. Lister427-Cas9-AnTat1.1 with VSG-228 experiments performed by K.J.W. Molecular cloning performed by J.E.S., K.J.W., E.M.K., and D.N.M. Transgenic parasites obtained by J.E.S., K.J.W., and E.M.K. Western blots performed by J.E.S. and E.M.K. VSG-AMP-seq pipeline written by J.E.S. RNA extractions, DNase treatment, and VSG PCRs for sequencing performed by J.E.S., K.J.W., and A.K.B. VSG-seq performed by J.E.S and A.K.B. VSG-AMP-seq performed by J.E.S. and K.J.W. Mouse infections and all downstream analysis performed by J.E.S. ORF mosaic recombination identification was performed by J.E.S. and A.M. Mosaic parasite clones processed for sequencing by J.E.S., K.J.W., and J.Z. FACS experiments and analysis performed by J.E.S. Modeling of VSGs performed by J.S. and L.M. Lister427 2018 VSG annotation pipeline written by J.E.S. VSG clustering performed by J.E.S., J.S., A.M., J.M.C.H., and E.H.A.

## Declaration of Interests

The authors declare no competing interests.

## Supplemental Information

Document S1. Figures S1-S5 Table1.

## Supplemental figures

**Supplemental Figure 1.**
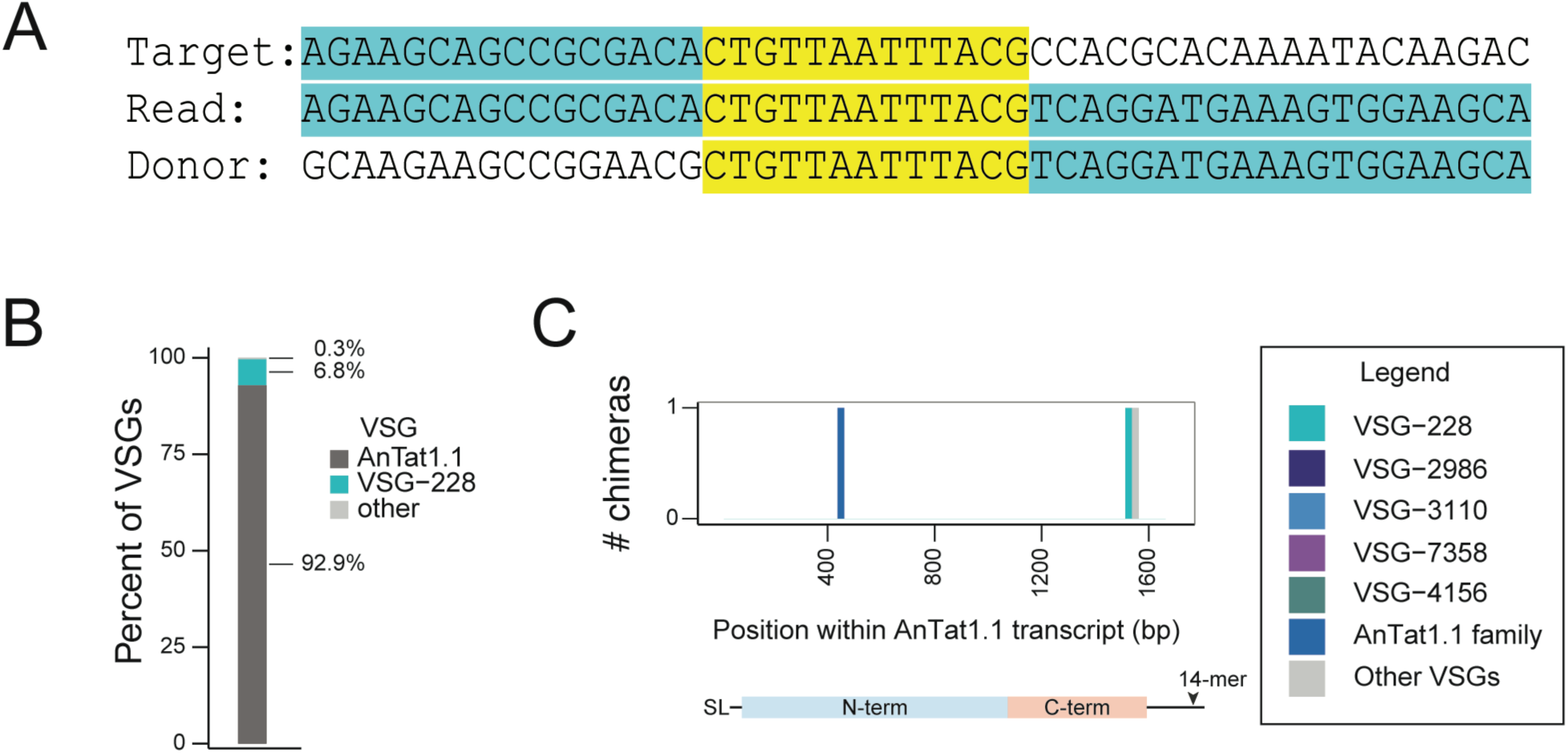
Mosaic Recombination Site & VSG-AMP-seq Validation. A) A schematic of a recombination site. The region in yellow is the putative recombination site, shared by all 3 sequences. B) A stacked bar graph quantifying the mix of Lister427 parasites engineered to express either AnTat1.1 or VSG-228 as determined by VSG-seq. These VSGs are naturally absent from Lister427 parasites and therefore chimeric reads containing sequence from both VSGs must have arisen from error. C) A histogram showing the midpoint of chimeric recombination events misidentified as a mosaic VSGs by VSG-AMP-seq from the mixed parasite sample in Supplemental Figure 1B.

**Supplemental Figure 2.**
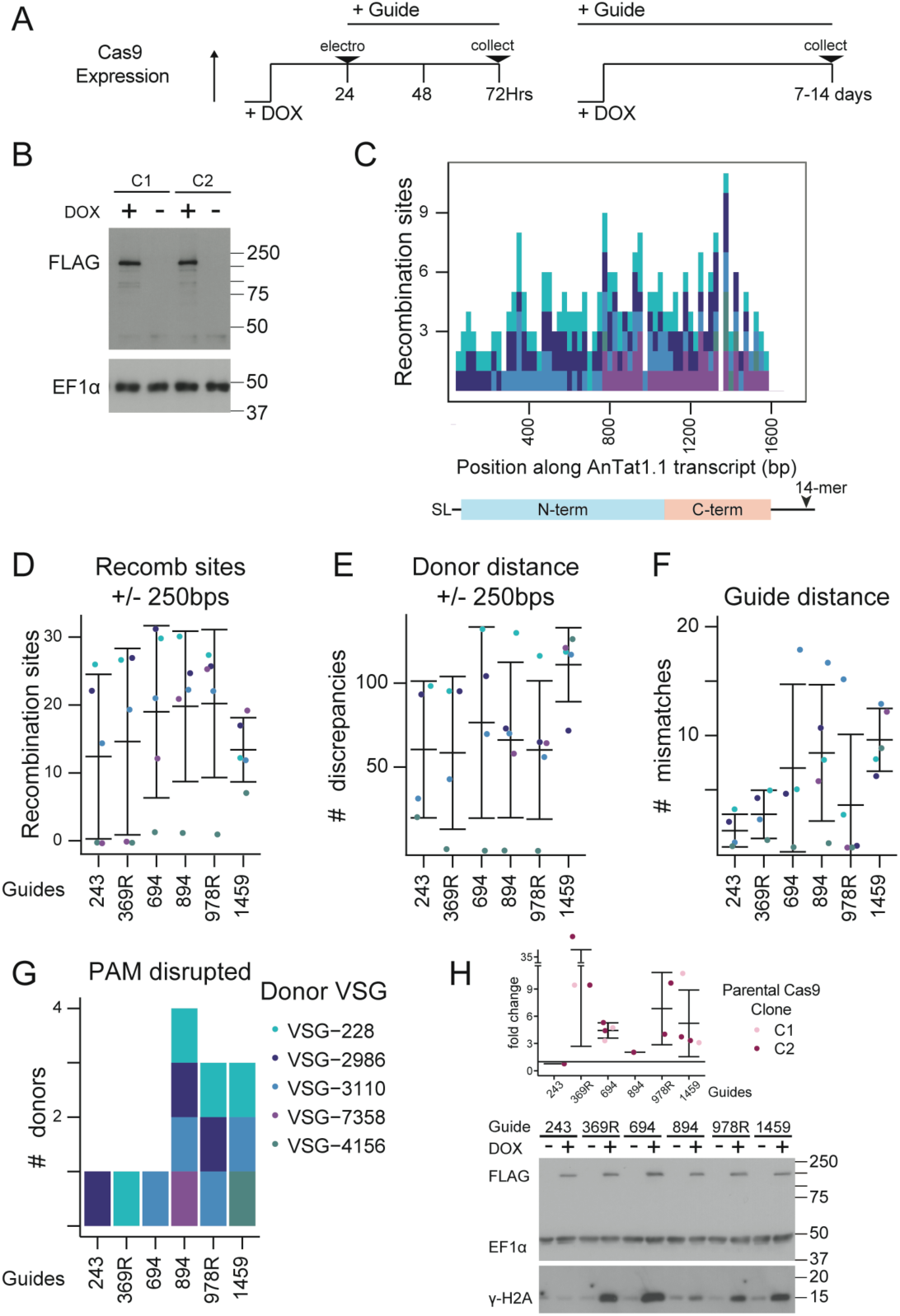
Cas9 system and sgRNA design and analysis. A) A schematic of the Cas9 induction experiment. Transient electroporation shown on the left while constitutive guide expression to isolate mosaic clones shown at right. B) An immunoblot showing FLAG-tagged Cas9 induction 24 hours following doxycycline treatment (DOX). DMSO was used as a vehicle control. EF1α was used as a loading control. C) A histogram showing the midpoint of all possible recombination sites 5bps or longer between AnTat1.1 and its family members. D) Quantification of the number of recombination sites within 250bps up or downstream of the cut site for each donor VSG. Mean ± s.d. E) Quantification of the Levenshtein distance between AnTat1.1 and family members. This includes mismatches, insertions and deletions. Mean ± s.d. F) Quantification of the number of mismatches at the guide binding site between AnTat1.1 and family members. Mean ± s.d. G) Histogram of which donor VSGs can disrupt the PAM when used to repair AnTat1.1. H) Quantification of the induction of DNA damage following guide induction as measured by a proxy of DNA damage, γ-H2A phosphorylation, following 24 hours of induction. γ-H2A induction was normalized to EF1α loading and fold change of staining intensity was determined between doxycycline-induced and uninduced samples. DMSO was used as a vehicle control. Mean ± s.d. Below, a representative immunoblot of FLAG, EF1α, and γ-H2A from doxycycline induced clones and uninduced controls is shown. Multiple independently-generated clones were induced. SL = 5′ splice leader sequence, 14-mer = 3′ sequence conserved in all VSG transcripts

**Supplemental Figure 3.**
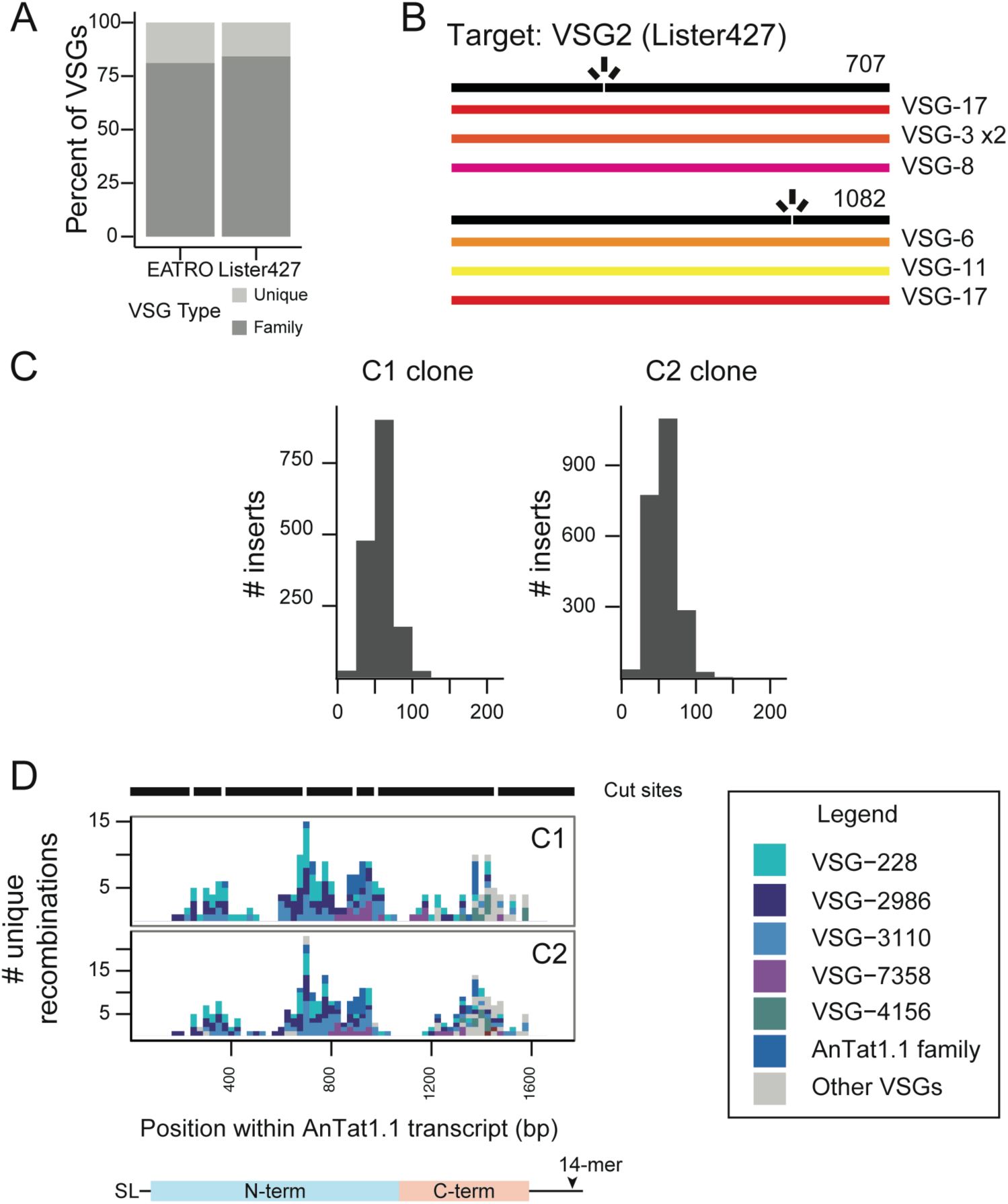
DNA breaks result in switchers if a homologous donor is not present. A) Quantification of the types of VSGs within the VSGnomes from EATRO1125 and Lister427 Parasites using the UCLUST greedy clustering algorithm. The Lister427 VSGnome has 4 VSGs which are perfectly duplicated without other family members. B) A schematic showing identified switchers following a Cas9-induced double strand break within VSG-2 at locations 707 and 1082. C) Histograms of donor VSG insertion lengths identified in all mosaic VSG reads from Figure 2A). The insert length only includes newly inserted sequence and does not include recombination sites. Clone C1 and C2 sequenced via VSG-AMP-seq are shown separately. The limit of detection for an insertion is approximately 200 bp. D) A histogram with a summary of the unique recombination sites found within guide-induced break for each clone. Cut sites are indicated above the histograms as gaps within the black line. The midpoint of the perfect homology between AnTat1.1 and the donor VSG at the recombination site is plotted. If a mosaic sequence matched >1 potential donor VSG, the average recombination position was plotted. The legend for the donor VSG colors is to the right. SL = 5′ splice leader sequence, 14-mer = 3′ sequence conserved in all VSG transcripts

**Supplemental Figure 4.**
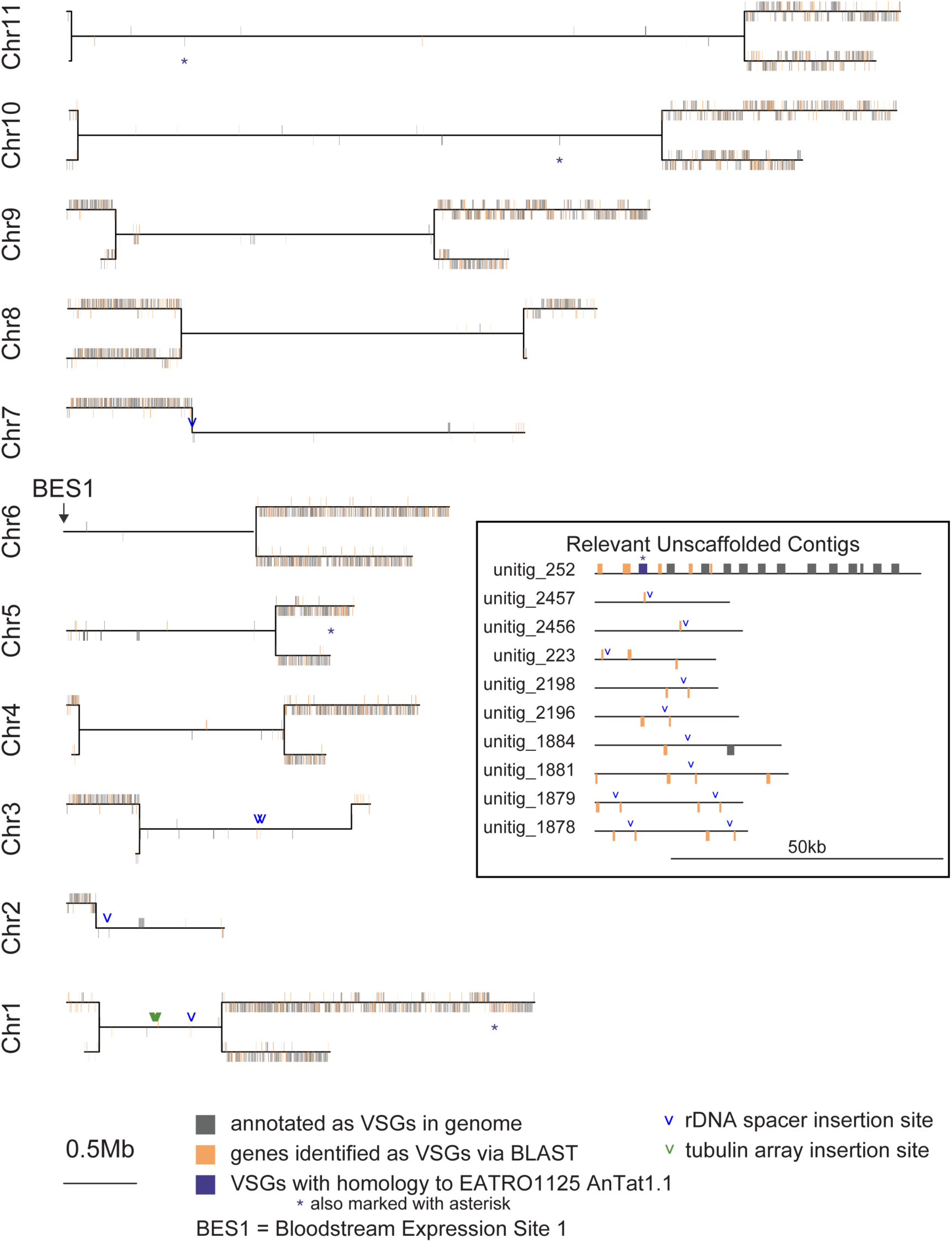
The Lister427 VSG annotated genome. The megabase chromosomes from Müller et al.^23^ VSGs annotated in the genome are plotted in gray. Unknown genes identified as VSGs via BLAST are plotted in yellow. Details of how these VSGs were identified are detailed in Methods. AnTat1.1 homologous family members are colored purple and marked with an asterisk. Insertion sites for the VSG-228 are denoted by arrows at the insertion location. Inset are the unitigs which are unscaffolded and harbor a copy of the AnTat1.1 family member or a potential insertion site. Bloodstream Expression Site 1 (BES1) is on the 5’ end of chromosome 6 and is marked by an arrow.

**Supplemental Figure 5.**
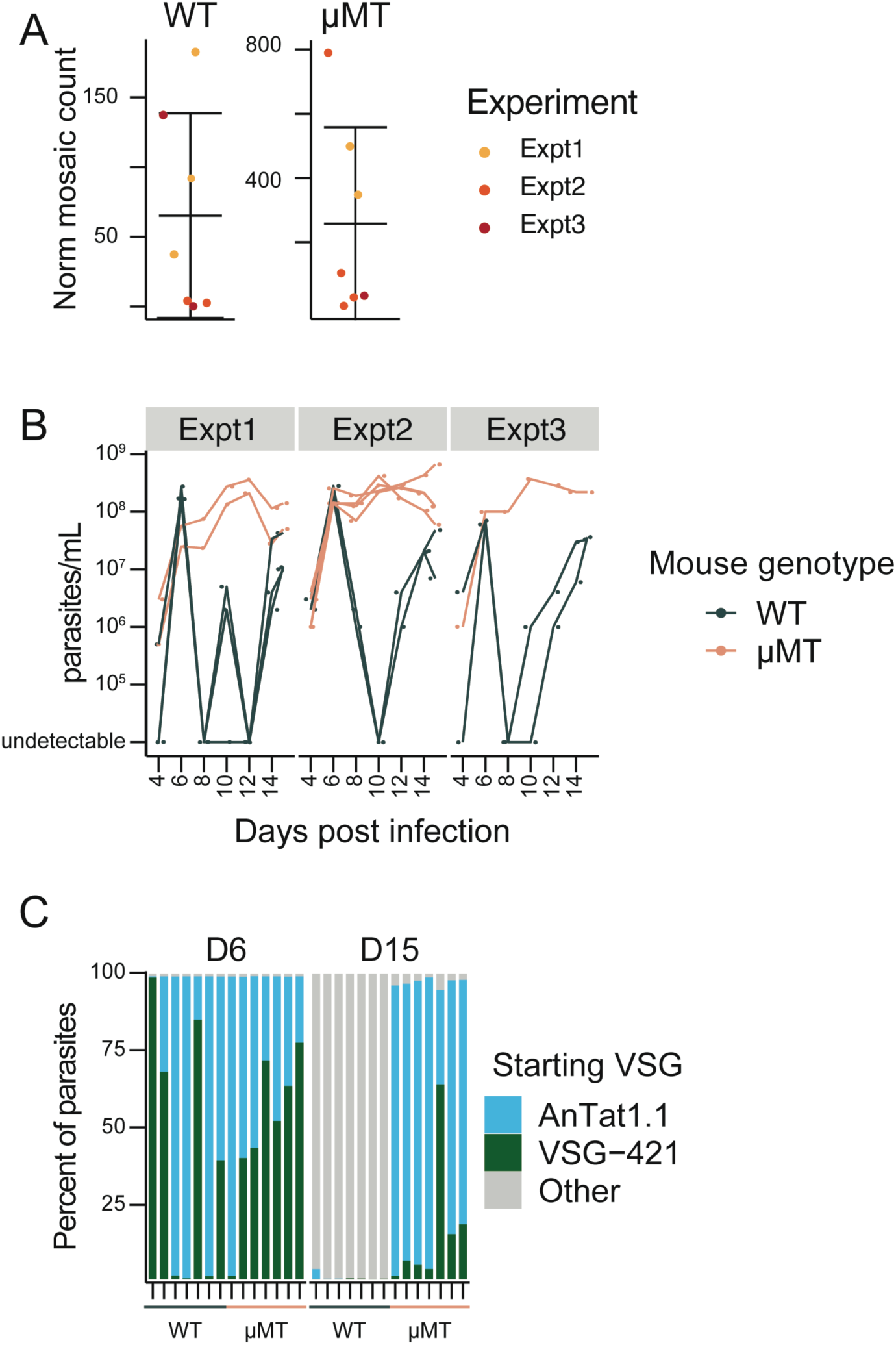
Mouse infection extended data. A) Quantification of the number of recombination events detected per mouse. Each mouse was normalized to the number of total consolidated, aligned, and unanchored reads compared to the consolidated, aligned, unanchored read count for one mouse to control for sequencing depth. This was performed separately for each genotype. Mean ± s.d. B) A time course of the parasitemia for the mouse infections. Blood was harvested every two days. C) Quantification of the percent of parasites expressing the starting VSG at D6 and D15 post-infection as quantified by VSG-seq.

## Notes

### Competing Interest Statement

The authors have declared no competing interest.

### Summary of Updates

Manuscript updates include: validation of VSG-AMP-seq with control samples (Supplemental Figure 1B & 1C); Additional mosaic clones obtained and Figure 2 updated (Figure 2C & 2D); updated analysis of VSG families (Figure 2G & Supplemental Figure 3A); Added an experiment to determine donor VSG accessibility within the genome (Figure 4); Updated discussion to reflect changes above; Expanded methods to include experiments above; Added author contributions section; Uploaded all associated data to SRA: PRJNA1140873; Code posted to: github.com/mugnierlab/Smith2024; Supplemental Figure 4 added to show positions of donor insertion in genome + expanded Lister427 VSG repertoire

